# REM sleep has minute-scale rhythms in mice and humans: A non-binary continuum between phasic and tonic microstates

**DOI:** 10.1101/2022.07.11.499612

**Authors:** Lezio S. Bueno-Junior, Maxwell S. Ruckstuhl, Miranda M. Lim, Brendon O. Watson

## Abstract

Rapid eye movement sleep (REM) is believed to have a binary temporal structure with “phasic” and “tonic” microstates, defined by motoric activity versus quiescence, respectively. However, we observed in mice that the frequency of theta activity (a marker of rodent REM) fluctuates in a non-binary fashion, with the extremes of that fluctuation correlating with phasic-type and tonic-type facial motricity. This demonstrates that phasic and tonic REM rather represent ends of a continuum. These cycles of brain physiology and facial movement occurred at 0.01-0.06 Hz, or infraslow frequencies, and affected cross-frequency coupling and neuronal assembly activity in the neocortex, suggesting network functional impact. We then confirmed that humans also demonstrate non-binary phasic/tonic microstates, with continuous 0.01-0.04 Hz respiratory rate cycles matching the incidence of eye movements. Thus, we discovered a fundamental property of REM, which can yield new insights into our understanding of sleep health.

## Introduction

Across mammals, rapid eye movement sleep (REM) is characterized by wake-like brain activity, atonia of postural muscles and myoclonic events, such as facial and limb twitches (Blumberg et al., 2020; Chase, 2013; Luppi and Fort, 2019). REM was formerly considered a uniform state, but is now believed to have at least some degree of sub-structure composed of dichotomous “phasic” and “tonic” periods, which respectively show high and low incidence of rapid movements (Simor et al., 2020). In humans, phasic REM is less susceptible to awakening and less sensitive to sensory stimuli, despite being motorically active (Ermis et al., 2010; Koroma et al., 2020; Sallinen et al., 1996; Takahara et al., 2002; Wehrle et al., 2007), whereas tonic REM is more reactive to stimuli and electrographically more similar to WAKE, despite being quiescent (Simor et al., 2018, 2019).

While understudied in rodents, phasic and tonic REM do seem present in such animals, with studies showing differences in oromandibular motricity (Brooks and Peever, 2008; Burgess et al., 2008; Morrison et al., 2003a, 2003b) and network activity within the hippocampus (Mizuseki et al., 2011; Montgomery et al., 2008). Although these kinds of studies have broad implications for sleep health and REM-related memory consolidation, they do not directly provide detail in distinguishing the REM microstates. A REM-focused study (Brankačk et al., 2012) contributed more directly to that distinction in mice. The authors found that brain oscillations in the theta band (5-10 Hz, typical of rodent REM) peak at higher frequencies in phasic REM (9-10 Hz) and lower frequencies in tonic REM (5-6 Hz), reinforcing the neurophysiologic heterogeneity within REM.

In addition to the paucity of studies on REM microstates in rodents, even in humans the methods for distinguishing these microstates are heterogeneous. There is no defined timescale to subdivide REM, with time bins ranging from 3 to 30 s (De Carli et al., 2016; Tamaki and Sasaki, 2019), nor is there a consensus on what criteria to use to categorize these time bins as phasic or tonic, given the diversity of sorting metrics (e.g., number of electrooculographic deflections; Koroma et al., 2020; Simor et al., 2016). The same inconsistency is observed in rodent studies, whose metrics to classify REM microstates vary from electromyogram to brain oscillations (e.g., Burgess et al., 2008; Mizuseki et al., 2011). And even if we had standardized methods to study phasic and tonic REM, simply comparing these microstates cannot elucidate the periodicity with which they alternate, or the extent to which they are dissociable at all.

Therefore, we still lack a fundamental understanding of REM microstructure. In this study, we address this knowledge gap in a comprehensive manner, at levels ranging from motor to brain activity in both mice and humans. First, we electrophysiologically recorded mouse REM structure both in the home-cage to enable natural sleep and under head fixation to provide fine detail on facial movements in conjunction with brain physiology. We then asked: do microstates alternate rhythmically throughout a REM episode? Are microstate transitions really unambiguous? Are there intra-REM fluctuations in brain activity, muscle tone and myoclonic bursting? Are these fluctuations related or independent? Based on what we learned from mice, we then explored a polysomnography dataset from healthy humans with a focus on fluctuating physiological signals suggestive of REM microstructure.

## Results

### Flexible freely-moving / head-fixed sleep recording and REM epoching in mice

We aimed to quantitatively determine the relationship between behavioral and electrophysiologic features of REM sleep. To do so, we established mouse recordings with three features: (1) flexible transferring of the same mouse between freely moving and head-fixed sleep setups (Figure 1A-B), (2) precise extraction of REM epochs (Figure 1C-D), and (3) laminar recordings from barrel cortex and hippocampus using a single stereotaxic implant (see Figure S1A-C and STAR Methods).

**Figure 1.**
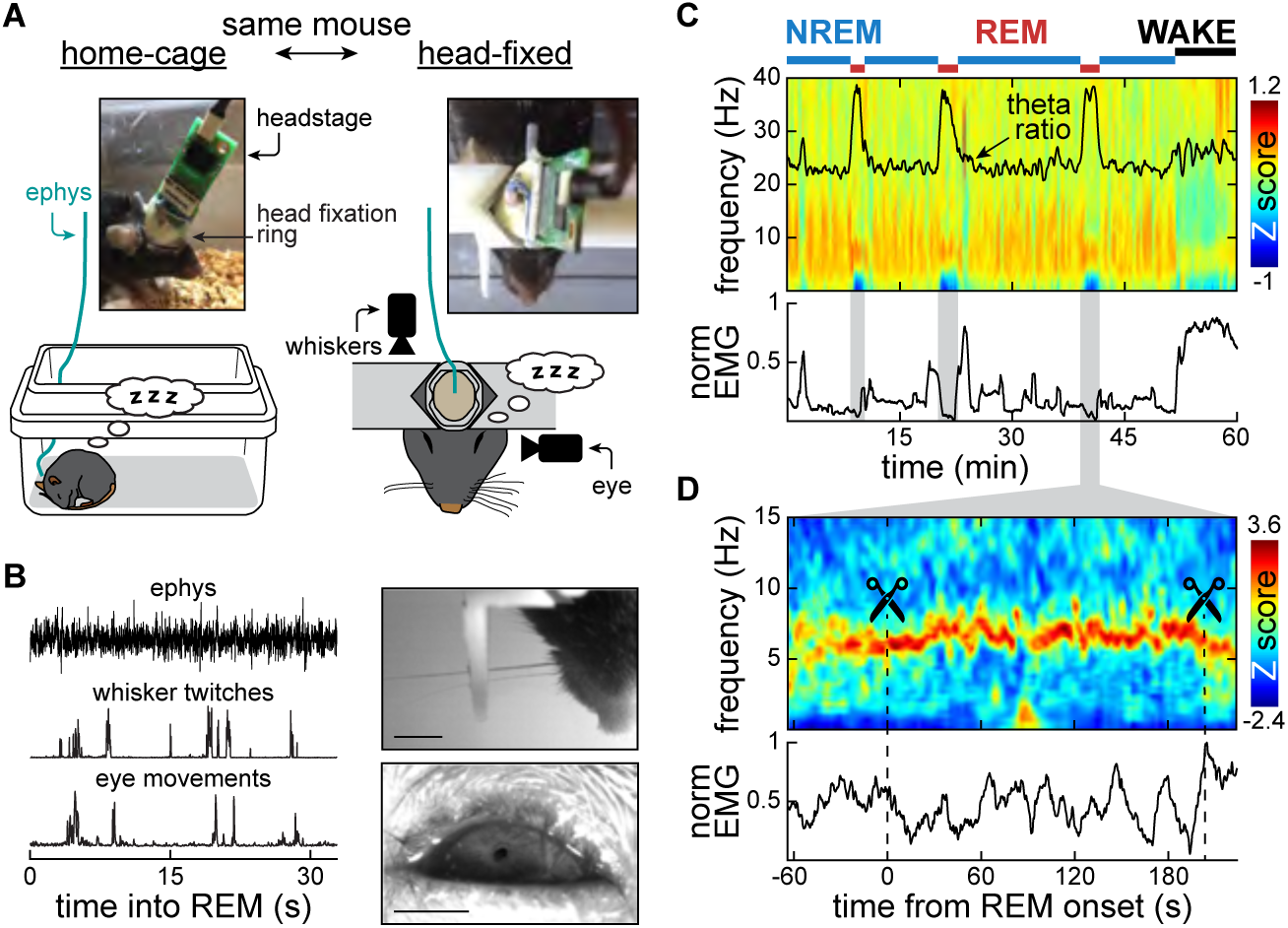
Flexible freely-moving / head-fixed sleep recording and REM epoching in mice. **(A)** Dual-mode chronic implant. Left drawing: freely moving electrophysiology. Left photo: cemented head cap embedding a 3D-printed RIVETS ring (Osborne and Dudman, 2014) and the silicon probe connector (Neuronexus). Plugged into the connector is the digitizing headstage (Intan) and its cable. Right drawing: 3D-printed RIVETS clamps forced against the ring-containing head cap for stable video and electrophysiology recordings from brain, whisker and eye during sleep. Right photo: same mouse and head cap system but head-fixed. For brain implantation, we used a linear silicon probe targeting both barrel cortex and hippocampus (Figure S1A-C). **(B)** Synchronized electrophysiology and videography to dissect the microstructure of REM (see also Video S1). Left: representative alignment of local field potentials (LFP) and rapid movements into REM-containing epochs. Top right: untrimmed whiskers with somatotopic correspondence to the implanted barrel cortical area (Figure S1A-C). Scale: 3 mm. Bottom right: eye imaging using infra-red reflectance thanks to the incomplete eyelid closure we typically observed during sleep (similar to Yüzgeç et al., 2018 and Sattler and Wehr, 2021). Scale: 1 mm. **(C)** Sleep scoring in 1-s bins (Grosmark et al., 2012; Watson et al., 2016). Bins were algorithmically scored as non-REM sleep (NREM), REM, or WAKE based on LFP and LFP-derived electromyogram (EMG; Schomburg, 2014). Scores were then manually adjusted using a state editing interface. REM epochs were extended in both directions to include 30–60-s margins around theta ratio peaks (gray areas). **(D)** REM margin trimming at higher temporal resolution. Wavelet spectrograms at 125-Hz sampling clarified REM boundaries, characterized by transitions to and from concentrated theta activity (5-10 Hz) in hippocampal channels. Refined epochs emphasized intra-REM fluctuations (see between dashed lines).

#### Dual-mode sleep recordings

Home-cage recordings have for years been used to study sleep in rodents, and we were able to repeat that here. In addition, habituated mice were able to reach REM under head fixation (Yüzgeç et al., 2018), allowing us to add high resolution videographic data on rapid eye and whisker movements to simultaneous intracranial recordings (Figure 1A-B; Video S1).

#### REM extraction and amount of REM episodes

REM epochs were extracted using a sleep scoring pipeline for rodents, in 1 s bins (Grosmark et al., 2012; Watson et al., 2016; Figure 1C). REM epochs were then refined at higher temporal resolution (125 Hz; Figure 1D). Both methods were based on local field potentials (LFP) and LFP-derived electromyogram (EMG; Schomburg, 2014). Resolving REM epochs at higher precision was important for this study, given our focus on REM microstructure. From six mice, we acquired a dataset with 312 home-cage and 76 head-fixed REM episodes. Although these numbers confirm that head-fixed REM is more difficult to achieve, the proportion of REM episodes across mice was relatively even between recording methods (Figure S1D). Figure S1D-E additionally shows details on the incidence and duration of REM episodes (comparable with Tsunematsu et al., 2020).

### Rapid movements tend to burst in discrete periods of mouse REM

First, we analyzed head-fixed videography to investigate whether facial twitching in mice supports the existence of two REM microstates. Figure 2A-B presents LFP and video data from a representative head-fixed REM episode. As displayed, saccading, eye blinking and whisking do have times of relative quiescence and others of bursting activity (Figure 2C), which at first glance confirms microstate-like epochs in REM.

**Figure 2.**
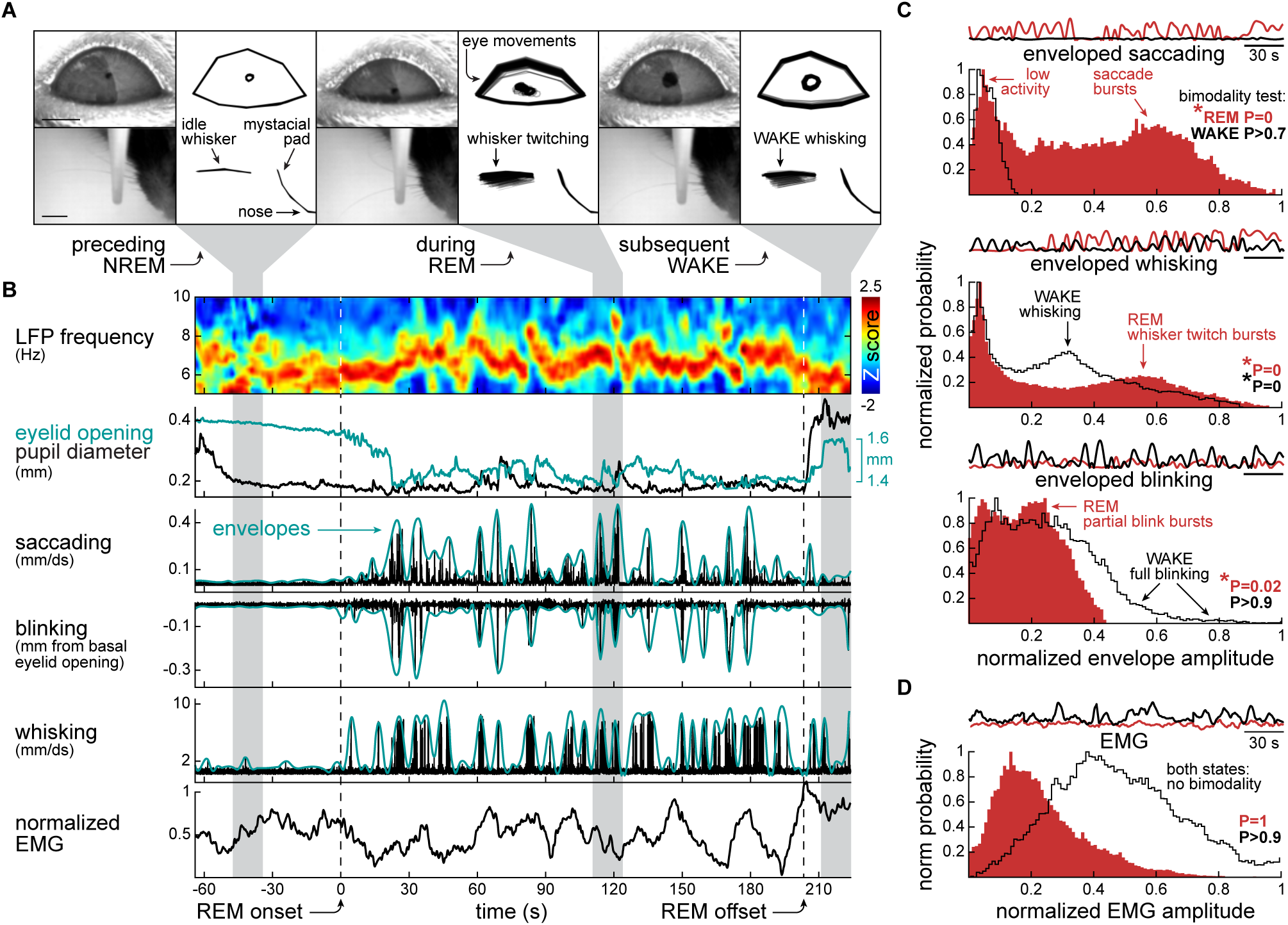
Rapid movements tend to burst in discrete periods of mouse REM. **(A)** Rapid eye and whisker movements were tracked using DeepLabCut (Nath et al., 2019) and custom code, resulting in simplified shapes and lines on the eyelids, pupil and whiskers. Three equally sized time intervals around a REM episode are illustrated (gray funnels). Deviations from basal positions appear in lighter tones. Scale bars: 1 mm (eye/pupil), 3 mm (whisker). See also Video S1, Figure S2 and Table S1. **(B)** Time-locked electrophysiology and videography. From top to bottom: theta-band wavelet spectrogram, eyelid opening (vertical distance between eyelids, followed by moving average), pupil diameter (horizontal width of the pupil), saccading (speed of the pupil in millimeters per decisecond, mm/ds), blinking (raw eyelid opening subtracted by its moving average), whisking (mean speed of all whisker labels in mm/ds) and EMG. Saccading, blinking and whisking were characterized by spike-like events. Such events often occurred in bursts, which we captured using peak envelopes (2-s window). **(C)** Distributions of rapid movement envelopes in REM vs. WAKE. Same-size REM-WAKE epoch pairs were jointly normalized, as represented by the curves at the top of each subplot. These illustrative curves do not contain all data, which are instead shown in the histograms below them (100 bins). Data were statistically examined for bimodality (Hartigan’s dip test), with test results shown as P values. All facial movement envelopes formed bimodal distributions in REM, suggesting alternation between REM sub-states. **(D)** Similar analysis but from EMG, which was predictably higher in WAKE than REM due to the postural atonia in REM. No bimodality was found in REM, unlike the analysis of facial twitching in (C).

#### Eye and whisker tracking

Figure 2A illustrates how eye and whisker movements were analyzed using DeepLabCut (Nath et al., 2019) and custom code. Notable is the incidence of movements in REM and WAKE, but not NREM. Additionally, partial eyelid opening and smaller pupil diameter were observed during sleep, replicating observations from prior studies in head-fixed (Tsunematsu et al., 2020; Yüzgeç et al., 2018) and freely moving mice (Sattler and Wehr, 2021). See also Video S1 to appreciate the movements in real time. See Figure S2 and Table S1 for additional details on our video dataset.

#### Enveloping of facial movement bursts

Figure 2B illustrates the alignment of a theta-band spectrogram and EMG tone with video-derived measures (see STAR methods). Here we focused on saccading, blinking and whisking, which exhibited spike-like events, similar to previous studies using non-videographic recording of rapid eye movements in rodents (Massot et al., 2019; Meng et al., 2021; Sánchez-López and Escudero, 2011, 2015). Relevant to intra-REM rhythms, facial movements occurred in bursts, which we captured using peak envelopes with 2 s window (Figure 2B).

Enveloped facial movements in REM were examined for bimodality of amplitude, with the hypothesis that periods with high and low motor activity would form two modes suggestive of phasic/tonic microstates. WAKE movements were used as controls. Thus, envelopes from REM and WAKE epochs were normalized together within each recording session for homogenization across mice (n = 6), resulting in Figure 2C. Each subplot contains representative envelope sub-epochs, and histograms from the whole dataset. Saccading and blinking were both bimodal in REM but not WAKE, indicating microstates of low and high oculomotor activity in REM. Whisking was bimodal in both REM and WAKE, but the motor activity mode of REM reached higher normalized values, reflecting the spasmodic kinects of REM whisking (Figure 2C). We also analyzed the distribution of EMG samples across the same normalized scale (Figure 2D) for comparison between facial movements and the muscle tone. No significant bimodality was found, and EMG was generally lower in REM. This is consistent with the well-known atonia of skeletal muscles in REM (Burgess et al., 2008; Chase, 2013).

Thus, enveloped facial movements in mouse REM do form bimodal distributions. This supports the existence of two REM microstates.

### Theta frequency fluctuates in an infraslow continuum during mouse REM

However, Figure 3 shows the novel finding that theta frequency in mouse REM fluctuates continuously without plateauing in discrete microstates. Interestingly, we found this fluctuation to occur in approximately 1 minute cycles (infraslow rhythm).

**Figure 3.**
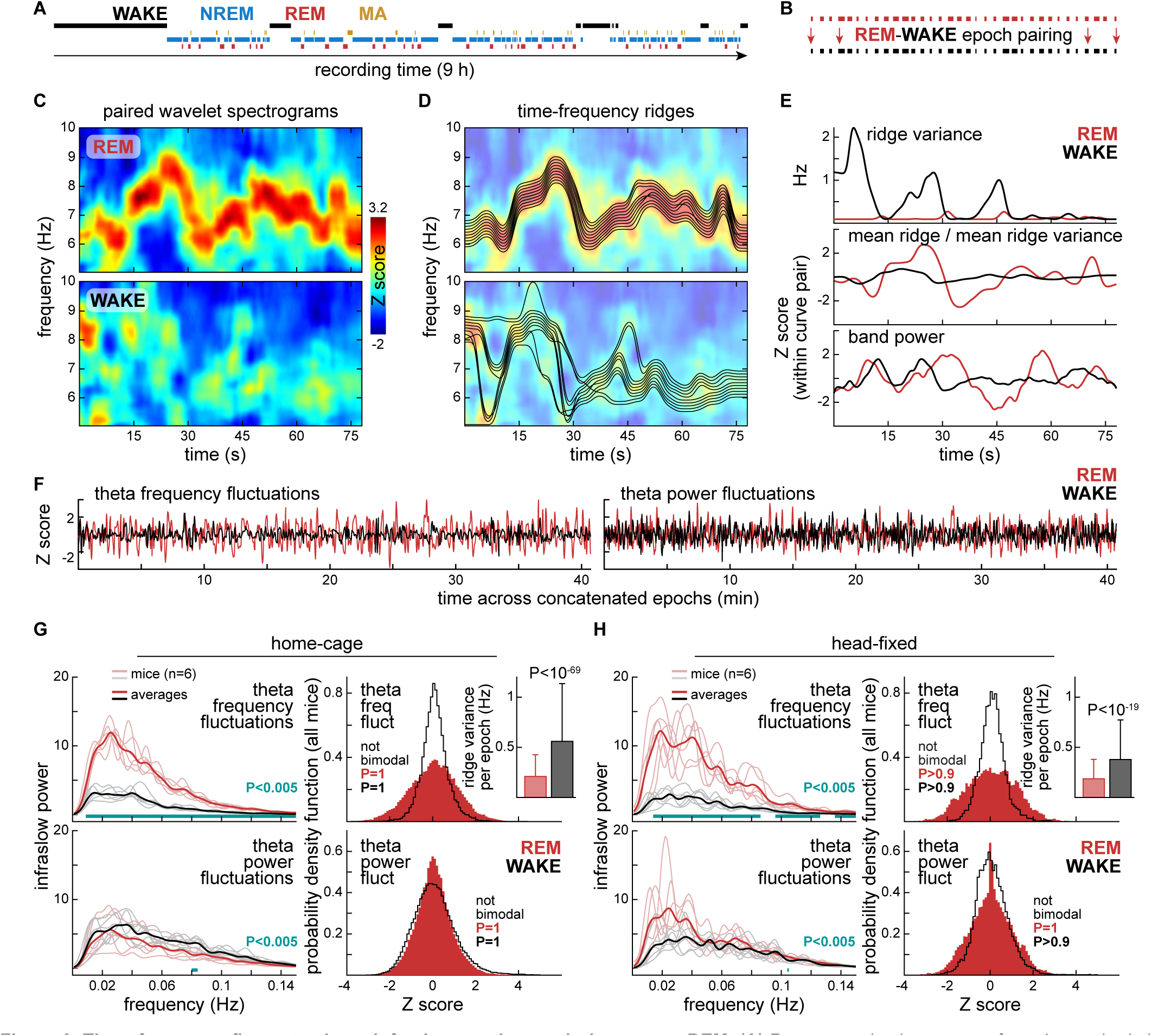
Theta frequency fluctuates in an infraslow continuum during mouse REM. **(A)** Representative hypnogram from the methods in Figure 1C. A full home-cage session is depicted. MA: microarousals. **(B)** For each post-trimming REM epoch (Figure 1D), we extracted a control epoch from the initial WAKE period of the same hypnogram. Control WAKE epochs were trimmed to the same number of time samples as their REM counterparts. WAKE epochs were separate from each other, without overlap of time samples. **(C)** Theta-band wavelet spectrograms from a representative REM–WAKE pair (125-Hz sampling). Spectrogram pairs were jointly Z scored along the frequency dimension. Theta activity was more organized in REM. **(D)** Same pair of spectrograms in (C) but overlaid with time-frequency ridges. The ten most dominant ridges were obtained per spectrogram. REM ridges were mostly adjacent to each other, finding contiguous power paths along time, unlike WAKE ridges, which showed greater variance. **(E)** Separate examination of theta frequency and theta power from the same spectrograms. Top: variance from the ten ridges in (D). Center: ridges were averaged and weighted (divided) by their mean variances. Weighted ridges were detrended and Z scored within each REM–WAKE pair. This resulted in stationary curves with cycles of several seconds. Bottom: theta power curves from averaging across the frequency bins of spectrograms, prior to their Z scoring. Theta power curves were also detrended and Z scored within each REM–WAKE pair. We again observed zero-centered waves lasting several seconds. **(F)** The curves explained in (E) were concatenated across REM or WAKE epochs, revealing minute-scale fluctuations. Theta frequency fluctuations showed higher amplitudes in REM. The time axes include all epochs represented in (B). **(G)** Spectral content and distribution of concatenated theta frequency and theta power fluctuations. Data from all home-cage sessions are shown (312 REM–WAKE epoch pairs, 6 mice; Figure S1D-E). Left: infraslow power spectral density (PSD) per mouse (faded lines) and averaged across mice (bold lines). Post-hoc differences at individual infraslow frequency bins are indicated (after two-way ANOVA). Infraslow rhythms were observed in both theta measures, with stronger PSD in REM theta frequency fluctuations. Right: distribution from infraslow fluctuations from all mice. We found no evidence of bimodality, suggesting that the infraslow oscillations are continuous, unlike the REM distributions in Figure 2C. The inset quantifies the mean ridge variance per epoch, explained in (E). Ridge variance was lower in REM, reflecting its organized theta activity. **(H)** As in (G), but from head-fixed data (76 REM–WAKE epoch pairs, 6 mice). Head fixation did not qualitatively change the physiology described in (G) except for slightly stronger infraslow PSD in theta power.

#### REM-WAKE epoch pairing, theta frequency and theta power

Figure 3A is a representative home-cage hypnogram from a mouse. Each REM epoch was paired with a duration-matched WAKE epoch from the same recording (Figure 3B). Figure 3C contains a representative REM-WAKE pair, shown as theta-band spectrograms from hippocampal LFP. We jointly Z scored each spectrogram pair, which resulted in frequency-varying undulating patterns, especially in REM (Figure 3C; like Brankačk et al., 2012; Karashima et al., 2005). These patterns were captured into numeric vectors using a time-frequency ridge algorithm. Figure 3D displays the behavior of these ridges, which were more contiguous in REM than WAKE (Figure 3D), leading to different ridge variances between the two states (Figure 3E, top). This implies that theta activity was more organized in REM than WAKE, so we compared the contiguity of multiple ridges within REM-WAKE spectrogram pairs. For that, ridges were separately averaged and weighted (divided) against their own variance. Averaged REM and WAKE ridges were then mean-subtracted for removal of slow trends and Z scored together. This resulted in zero-centered fluctuations in the scale of minutes, with stronger amplitudes in REM (Figure 3E, center). We then separately measured theta power. Spectrograms were averaged vertically resulting in band power curves, which were also detrended and Z scored. We again observed minute-scale fluctuations (Figure 3E, bottom). See Figure S3A-B for additional examples.

#### Infraslow rhythm and lack of bimodality in both theta activity measures

The Z-scored curves of Figure 3E suggested a predominant rhythm at frequencies less than 0.1 Hz, i.e., infraslow band (Lecci et al., 2017; Mitra et al., 2018; Osorio-Forero et al., 2021; Watson, 2018). To quantify this, we performed a spectral analysis using a minute-scale sliding window. To provide enough oscillatory data for that window to slide along, we concatenated Z scored curves across REM or WAKE epochs per recording session, as illustrated in Figure 3F. We then constructed power spectral density (PSD) plots (Figure 3G-H) showing the entire home-cage and head-fixed datasets, respectively. REM showed stronger infraslow PSD of theta frequency fluctuations in both home-cage (interaction: F_1,10_ = 495.14, P = 7.54*10^−10^) and head-fixed recordings (interaction: F_1,10_ = 353.73, P = 3.91*10^−9^), according to two-way ANOVA with frequency bins as repeated measures. The difference in the infraslow PSD of theta power fluctuations was less significant between REM and WAKE (home-cage interaction: F_1,10_ = 5.11, P = 0.047; head-fixed interaction: F_1,10_ = 7.42, P = 0.021).

We then interrogated whether theta activity measures vary continuously or are distributed in a bimodal manner. Evidence of bimodality would match our previous observations on facial movements (Figure 2C), and corroborate the concept of phasic and tonic microstates in REM. However, we found no significant bimodality, irrespective of brain state, theta activity measure, or type of recording (Figure 3G-H). In addition, theta frequency distributions showed higher variances in REM than WAKE, as shown in Figure 3G-H, top histograms (F-test; home-cage: F = 3.41, P = 0; head-fixed: F = 4.12, P = 0), consistent with the higher amplitude of REM theta frequency fluctuations (Figure 3F). Theta power distributions, in turn, were more similar between REM and WAKE, though still with significant differences in their variances (home-cage: F = 0.72, P = 0; head-fixed: F = 1.46, P = 0; Figure 3G-H). Lastly, the insets in Figure 3G-H quantify the mean ridge variance per epoch, i.e., the values used to weight ridge contiguity. Per-epoch ridge variance was lower in REM, both in home-cage (rank sum: P = 2.21*10^−70^) and head-fixed recordings (P = 6*10^−20^), reflecting the more uniform theta activity in REM.

By analyzing EMG as described above (Figure S3C-E), we also observed power fluctuations in the infraslow range, but with stronger PSD in WAKE than REM. This was true for both home-cage (Figure S3D; interaction: F_1,10_ = 650.09, P = 1.98*10^−10^) and head-fixed recordings (Figure S3E; interaction: F_1,10_ = 71.95, P = 7.03*10^−6^). Similar to theta activity measures, we found no evidence of bimodality (Figure S3D-E). Furthermore, WAKE EMG fluctuations exhibited greater variance in WAKE than REM (home-cage: F = 0.18, P = 0; head-fixed: F = 0.24, P = 0; Figure S3D-E). This suggests that WAKE EMG and REM theta frequency are comparable in terms of infraslow rhythmicity. Finally, as a control for our analysis of concatenated epochs, we examined infraslow PSD from individual epochs lasting more than 60 s, i.e., long enough for infraslow PSD without concatenation (Figure S3F-G). Results reinforce that REM shows stronger infraslow fluctuations of theta frequency (home-cage interaction: F_1,10_ = 159.31, P = 6.46*10^−30^; head-fixed interaction: F_1,10_ = 142.21, P = 2.27*10^−20^).

Therefore, theta frequency variations during REM may be a manifestation of infraslow brain rhythms, which we believe to entail fluctuations rather than bimodal flipping. This stands in contrast to the binary facial movement data. We address this paradox below.

### REM facial movements in mice preferentially accompany theta frequency fluctuations

#### Co-examination of video and electrophysiology variables in time bins

Figure 4A-C examines the temporal relationships between facial and electrophysiology variables, after binning both to 2 s. A summary is conveyed by correlation matrices (Figure 4A). In REM, we observed that the three types of enveloped facial movements were related to each other and to theta frequency, but not with theta power or EMG. The same relationships were observed in WAKE, in addition to EMG correlations.

**Figure 4.**
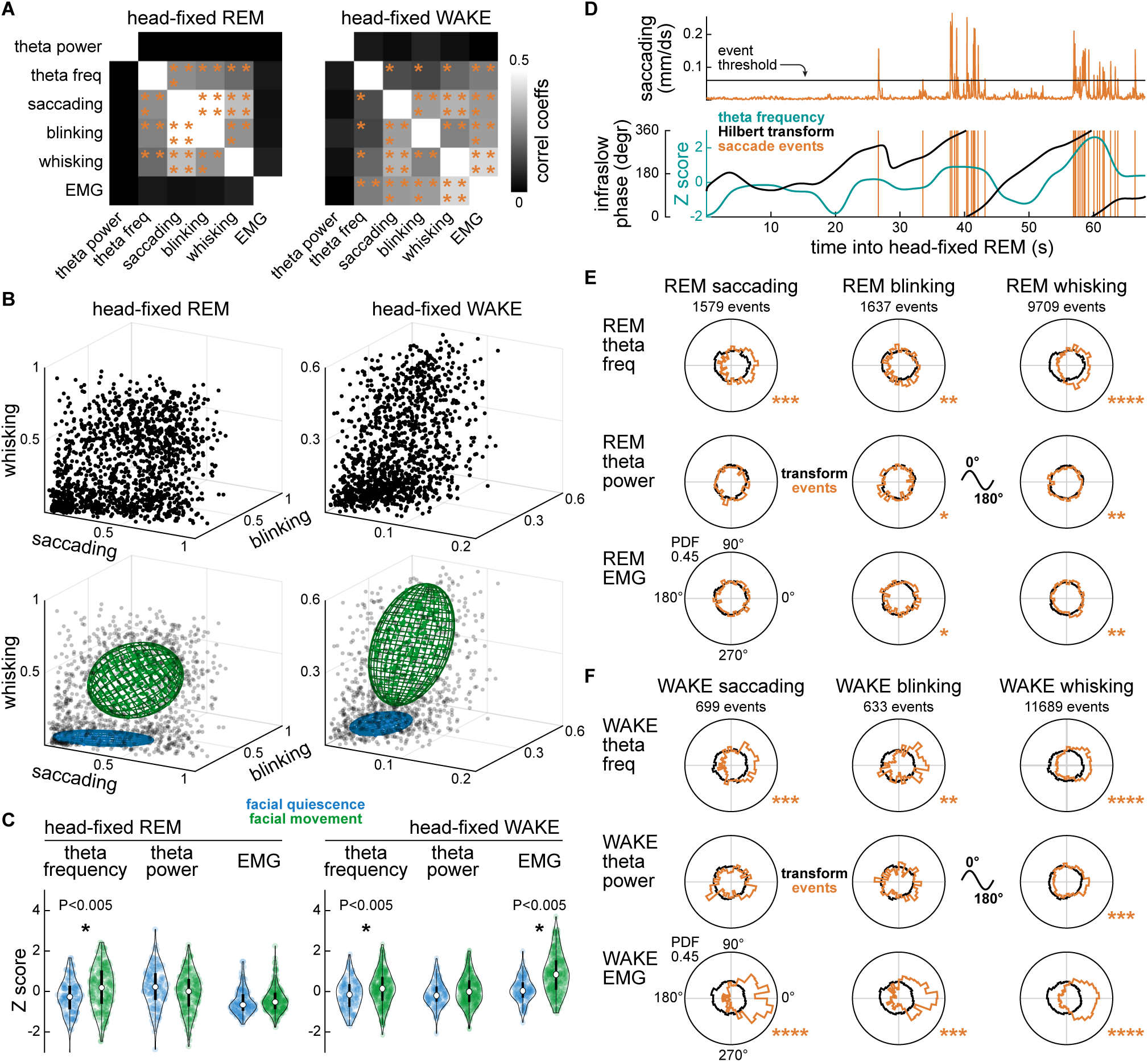
REM facial movements in mice preferentially accompany theta frequency fluctuations. **(A)** Matrices summarizing how enveloped facial movements and electrophysiological measures correlate. Variables from the head-fixed dataset were divided in 2-s bins, then examined for pairwise correlations (*P < 0.0017, **P < 10^−15^, ***P < 10^−27^, ****P < 10^−34^). In REM, positive correlations were confined to: (1) theta frequency with facial movement variables, and (2) facial movement variables with each other. Similar effects were found in WAKE, but along with EMG correlations. **(B)** Same time bins as in (A) scattered in a 3D space defined by saccading, blinking and whisking. By focusing on these dimensions, we explored if facial movements alone delimit REM sub-states. Top: facial movement variables formed overlapping clusters. Bottom: ellipsoids from a Gaussian mixture model illustrate data points belonging to facial quiescence or facial movement clusters. **(C)** Electrophysiological distinction between the clusters. P values at individual comparisons are post-hoc differences after two-way ANOVA. Left: during REM, the facial movement cluster showed higher Z scores in theta frequency. Right: during WAKE, the facial movement cluster showed higher Z scores in both theta frequency and EMG. Results are consistent with (A). **(D)** Facial movement events in relation to the phase of infraslow fluctuations. Top: raw, not enveloped, facial movement curves were upsampled from their frame rates (20 Hz in eye videos, 100 Hz in whisker videos) to the LFP/EMG sampling rate (125 Hz) and thresholded (dashed line, 2x standard deviation). Threshold crossings were taken as timestamps. Bottom: electrophysiological measures were Hilbert-transformed to obtain their infraslow phases. Facial movement timestamps were then quantified for phase preference in (E-F). **(E)** Polar histograms from the entire head-fixed dataset showing phase relationships between REM facial movements (columns of subplots) and electrophysiological signals (rows of subplots). Each histogram (normalized radius limit: 0.45; probability density function) shows the phase distributions of facial movement events (beige) on top of the phase distributions from the underlying Hilbert transforms (black). The latter vary slightly due to the exclusion of episodes with problematic videos (Table S1). Rayleigh tests: *P < 0.0028, **P < 10^−8^, ***P < 10^−25^, ****P < 10^−66^. We found that REM facial movements prefer the peaks of theta frequency fluctuations (i.e., faster theta), consistent with (A-C). **(F)** As in (E), but from WAKE. In this state, facial movement events showed preference for the peaks of both EMG and theta frequency cycles, also corroborating (A-C).

The same time bins were analyzed with a clustering approach. The top plots of Figure 4B display such bins in the 3D space formed by enveloped facial movement variables. We observed partially overlapping clusters, which we captured into non-overlapping clusters named “facial quiescence” or “facial movement” using a gaussian mixture model (Figure 4B, bottom). The clusters were then examined for electrophysiological measures (Figure 4C). In REM, the facial movement cluster showed significantly higher theta frequency (cluster effect: F_1,352_ = 72.62, P = 4.67*10^−16^). In WAKE, both theta frequency and EMG were higher in the facial movement cluster (cluster effect: F_1,393_ = 59.03, P = 1.25*10^−13^; interaction effect: F_1,393_ = 76.28, P = 7.17*10^−17^). These observations are consistent with Figure 4A.

#### Phase relationships between facial movements and infraslow cycles

In Figure 4D-F we explored whether facial movements prefer specific phases of the underlying infraslow fluctuations. For that, the curves from facial movements, i.e., saccading, blinking and whisking, were upsampled from their original rates to the theta/EMG rate (125 Hz). Facial movements were then thresholded (Figure 4D top; like Lerma and García-Austt, 1985). In turn, the infraslow fluctuations, which we hypothesized to be “infraslow clocks” for the facial movements, were Hilbert-transformed. Phases of movement events were then gathered from this transform (Figure 4D, bottom). Figure 4E-F polar plots show that in REM, facial movements showed strongest preference for theta frequency (Figure 4E, top row). The same was found in WAKE, which additionally showed phase relationships between facial movements and EMG (Figure 4F, top and bottom rows). Distributions involving theta power were generally more uniform. All of these phase relationships were aligned with the peaks of infraslow cycles (zero degrees). Thus, according to Figure 4E-F, facial movements are more likely to occur in periods of faster theta frequency (both REM and WAKE) and higher EMG (WAKE only), corroborating Figure 4A-C.

Therefore, facial twitching is more likely as REM theta reaches higher frequencies. This reconciles our two contradictory results of bimodally-distributed facial movements (Figure 2) and unimodally-distributed infraslow rhythm of theta frequency fluctuations (Figure 3).

### Laminar LFP accompanies the infraslow fluctuation of theta frequency in mouse REM

We next performed spectral analyses from all 64 channels of our linear probes, which were implanted across primary somatosensory cortex and hippocampus. Our aim was to explore neocortical-hippocampal laminar LFP in relation to intra-REM fluctuations.

#### Dividing laminar LFP into infraslow sub-epochs

Figure 5 focuses on the infraslow fluctuations of theta frequency, given its correlation with facial movements shown above. As illustrated in Figure 5A, these fluctuations were coarsened into 45° steps, and those were used to subdivide the raw multi-channel LFP into eight “infraslow categories”. These categories were then examined for spectral patterns at frequencies greater than 4 Hz (Figure 5A). Based on our findings described thus far, we interpreted the 0° and 180° categories as the “poles” of mouse REM (Figure 5B): the “phasic pole” (fast theta, facial movement) and the “tonic pole” (slow theta, facial quiescence), respectively.

**Figure 5.**
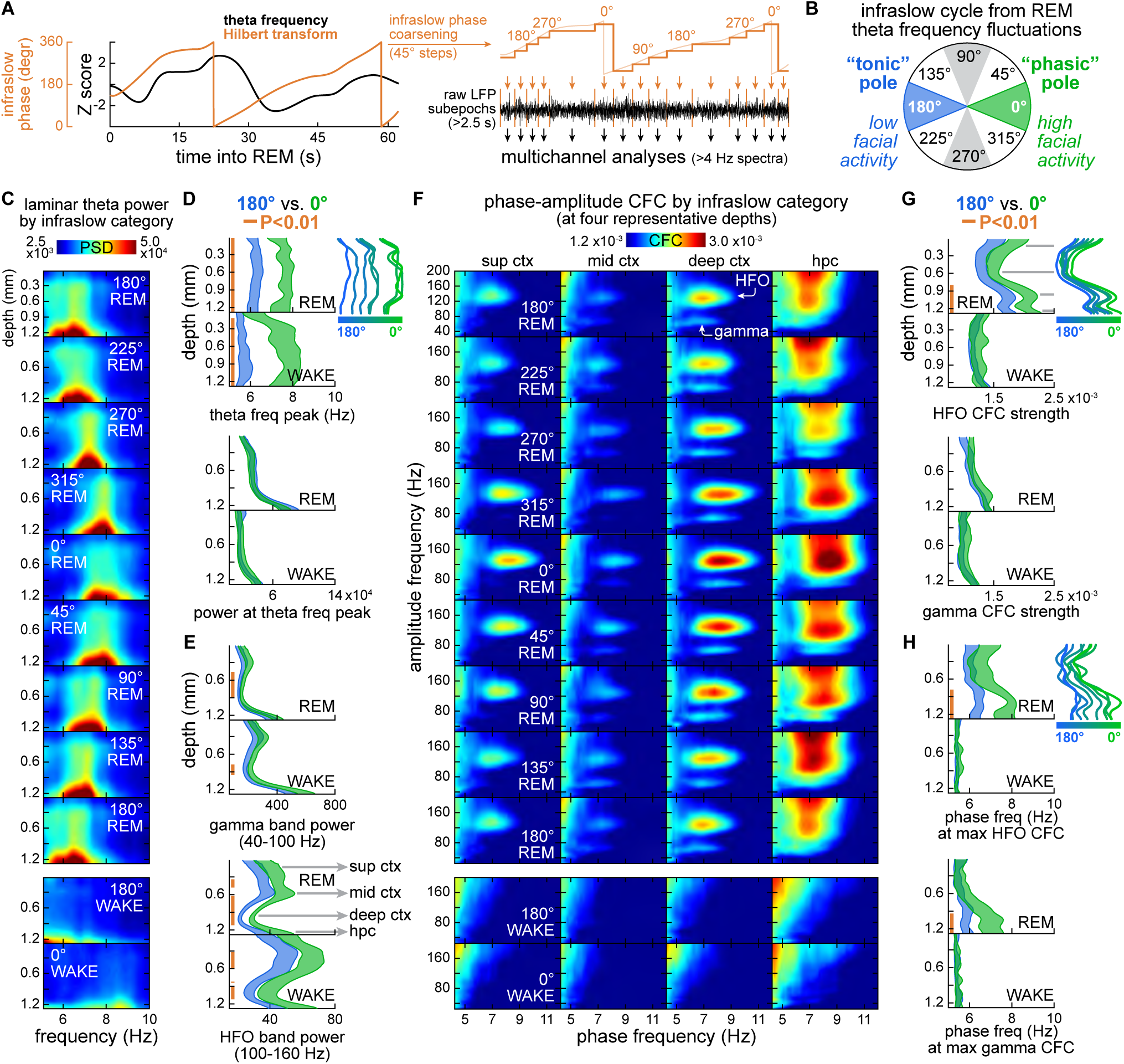
Laminar LFP accompanies the infraslow fluctuation of theta frequency in mouse REM. **(A)** Left: theta frequency fluctuations were Hilbert-transformed into infraslow phases, as in Figure 4D. Right: phases were coarsened into 45° steps, resulting in eight angle categories per infraslow cycle. Boundaries between 45° steps were used to delimit sub-epochs in the raw LFP. Sub-epochs longer than 2.5 s were examined for spectral patterns above 4 Hz. This procedure was iterated over 64 channels, constructing laminar profiles encompassing S1 cortex and hippocampus. **(B)** Infraslow phase categories from the fluctuations in REM theta frequency. All eight categories are represented by the heatmaps in (C) and (F). Categories centered on 0° and 180° were quantitatively compared in (D, E, G, H). 0° and 180° were respectively interpreted as the “phasic” and “tonic” ends of the infraslow continuum of theta frequency fluctuations given the results in Figures 3-4. **(C)** Laminar LFP by the coarse infraslow phases of theta frequency fluctuations. Power spectral density curves (PSD, color axis) along theta frequencies (5-10 Hz, X axes) were stacked across channels (Y axes), then averaged across mice (home-cage sessions only), constructing one laminar profile per infraslow category (vertical sequence of subplots). A continuity is observed between fast theta (0°) and slow theta (180°) in REM, reinforcing the findings in Figure 3. Strong theta power at the bottom of each subplot is of hippocampal origin. WAKE showed weaker laminar theta irrespective of infraslow category (only 0° and 180° from WAKE are shown, for simplicity). See also Video S2. **(D)** 0° vs. 180° comparisons within REM or WAKE. The top and bottom pairs of graphs respectively show theta frequency peak and power at theta frequency peak. Shaded curves delimit standard errors from across home-cage sessions. Vertical beige lines indicate post-hoc differences at individual depths after two-way ANOVA. 0° vs. 180° differences are seen in theta frequency but not theta power. The top-right inset shows average theta frequency peak profiles per infraslow phase, further illustrating the continuity between 0° and 180°. See also Figure S4. **(E)** As in (D) but showing laminar power at higher frequency bands. The top and bottom pairs of graphs respectively show 40-100 Hz power (gamma oscillations) and 100-160 Hz power (high-frequency oscillations, HFO). Both bands differed between 0° and 180° in both REM and WAKE, especially laminar HFO. HFO also showed clearer stratification. Horizontal arrows indicate four representative depths: superficial, mid and deep S1 cortex and hippocampus. These depths were analyzed in (F). **(F)** Cross-frequency coupling (CFC) between theta (phase frequencies: 4-12 Hz) and faster oscillations (amplitude frequencies: 20-200 Hz) per infraslow category from theta frequency fluctuations at the representative depths indicated in (E). HFO coupling was found to oscillate continuously between 0° and 180° in terms of both CFC strength (color axis) and theta phase (X axes). Gamma coupling was observed to fluctuate in theta phase only. These effects were more evident in deep S1 cortex and hippocampus. CFC was much weaker in WAKE regardless of depth or infraslow category. Comparable with Brankačk et al. (2012) and Scheffer-Teixeira and Tort (2017). **(G)** Similar to (D) but showing the full laminar profiles of CFC strength. 0° vs. 180° differences are seen only in HFO CFC, especially at deeper channels. This effect was specific to REM, unlike the state independence in (E). The top-right inset shows average laminar profiles of HFO CFC per infraslow phase, illustrating that in this measure the continuity between 0° to 180° is exclusive to REM. **(H)** As in (G) but showing the full laminar profiles of CFC strength peaks along the phase frequency dimension. 0° vs. 180° differences were specific to REM and present in both HFO and gamma. 0° vs. 180° differences were again concentrated at deeper channels in REM.

#### Laminar profile of theta activity along REM infraslow categories

Figure 5C shows laminar profiles (64 channels, Y axes) of PSD (color axis) in the theta range (X axes) by the eight infraslow categories (vertical sequence of subplots). Only home-cage data are shown, displaying average images across mice. In Figure 5C, we see theta peak frequencies oscillating smoothly between 5 and 10 Hz, expanding upon our earlier findings on infraslow continuity. A gradient along the spatial dimension is also observed, with strong power at hippocampal depths (bottom-most sites) to intermediate power in S1 cortical depths (Figure 5C). The zone of intermediate-power theta extending up into S1 cortical depths may be partially due to volume conduction emanating from hippocampal theta during REM. However, neocortical generation of theta also cannot be ruled out, given the non-diminution of theta power across much of the middle cortical depth. WAKE showed a different pattern, with weaker theta power across depths (Figure 5C, bottom). Only 0° and 180° infraslow categories of WAKE are shown for simplicity (see also Video S2).

Focusing on the extreme infraslow categories 0° and 180°, we analyzed theta peak frequency and theta power separately across the laminar profile (Figure 5D). Shaded curves represent the variation from all home-cage sessions. Significant differences in theta peak frequency were found across the depth profile, both in REM (F_1,28_ = 53.21, P = 6.09*10^−8^) and WAKE (F_1,28_ = 45.82, P = 2.37*10^−7^; Figure 5D, top). This was expected, as infraslow categories were obtained from theta frequency fluctuations. However, laminar theta power was statistically not different between 0° and 180° in either state (Figure 5D, bottom). Thus, laminar theta frequency and theta power do not necessarily co-fluctuate in REM, consistent with Figures 3 and 4. The inset on the top right of Figure 5D displays average laminar profiles of theta peak frequency from all infraslow categories.

#### Laminar profile of faster oscillatory activities along REM infraslow categories

The bottom graphs of Figure 5D suggest stratification in theta power. To explore stratification, we expanded our laminar PSD analysis to faster - and possibly more local - oscillatory bands, namely gamma (40-100 Hz) and high-frequency oscillations (HFO; 100-160 Hz). As shown in Figure 5E, we indeed found anatomically localized differences between the extreme infraslow categories, but they were present both in REM (gamma: F_1,28_ = 6.81, P = 0.014; HFO: F_1,28_ = 38.12, P = 1.15*10^−6^) and WAKE (gamma: F_1,28_ = 10.39, P = 0.003; HFO: F_1,28_ = 12.08, P = 0.002).

We then looked further for state-dependent stratification by co-examining theta and gamma/HFO activities using the phase-amplitude cross-frequency coupling (CFC) method (e.g., Brankačk et al., 2012; Scheffer-Teixeira and Tort, 2017). First, we narrowed our search down to four depths representing the superficial, intermediate and deep S1 cortex, and hippocampus, based on our prior mapping of HFO power (Figure 5E, bottom). We then analyzed CFC with phase frequencies ranging from 4 to 12 Hz (containing theta) and amplitude frequencies ranging from 20 to 200 Hz (containing gamma and HFO). CFC was repeated per selected depth and per 45° infraslow category. As shown in Figure 5F and Video S2, this measure revealed higher state dependencies, with gamma and HFO coupling being stronger in REM. Another pattern of Figure 5F is that HFO blobs fluctuate smoothly both within the phase frequency dimension (X axes) and the CFC dimension (color axis), with higher values occurring around the 0° infraslow category. Figure 5F also shows the heterogeneity of CFC across depths, similar to a comparable study in rats (Scheffer-Teixeira and Tort, 2017). These patterns were not seen in WAKE (Figure 5F, bottom plots).

We then quantified statistical differences in these CFC data. Figure 5G shows that 0° vs. 180° differences in theta coupling strength were confined to HFO during REM (F_1,28_=23.03, P=4.79*10^−5^). This is in contrast with HFO power, where 0° vs. 180° differences were present in both REM and WAKE (Figure 5E, bottom). Additionally, at deep cortical layers, we found that HFO CFC is strong despite locally weak HFO power, suggesting that the efficacy with which REM theta modulates HFO varies with the depth. Interestingly, this physiology-anatomy relationship was also modulated within REM, since HFO CFC in deeper channels was found to increase further at the “phasic pole” of REM (see post-hoc differences in Figure G). Moreover, similar to the infraslow fluctuations in theta frequency - and probably because of them - laminar HFO CFC was also found to fluctuate smoothly between the 0° and 180° poles (inset of Figure 5G). Finally, Figure 5H confirms that the position of HFO and gamma CFC blobs along the phase frequency axis shifts between slow and fast theta as the infraslow cycles recur. This phenomenon was exclusive to REM, and again observed to be heterogeneous across depths, with the bottom third of the depth profile accompanying the infraslow fluctuation more closely (HFO: F_1,28_ = 25.03, P = 2.75*10^−5^; gamma: F_1,28_ = 13.24, P = 0.001).

As an additional control for Figure 5, Figure S4 describes how infraslow fluctuations of theta power and EMG (rather than theta frequency) affect the laminar power and CFC outcomes. Overall, the smooth shifting described in Figure 5 is disrupted when theta power or EMG fluctuations are used to dictate the infraslow sub-epoching (Figure S4; see legend for further explanation). This reinforces that theta frequency is an optimal measure for dissecting REM microstructure.

Therefore, the infraslow fluctuation of theta frequency during REM drives uneven spectral patterns across neocortical-hippocampal depths, with the “phasic pole” of REM enabling more efficient theta-HFO coupling at deeper channels.

### Laminar firing and spike coordination also accompany the infraslow microstructure of mouse REM

#### Cortical spiking increases in REM periods with fast theta frequency

How is the spiking of individual neurons modulated by this REM infraslow rhythm? Figure 6A displays a depth vs. time raster plot from a representative REM period, with spikes (dots) shown along with Hilbert-transformed theta frequency fluctuations (phase curve). As illustrated in Figure 6B, for each unit we calculated a preferred infraslow phase (positions on X axis) and mean resultant length, or MRL (circle diameters). By scattering the resulting circles across the depth vs. infraslow phase space, we revealed patterns such as Figure 6C, which were quantified across animals in Figure 6D-E.

**Figure 6.**
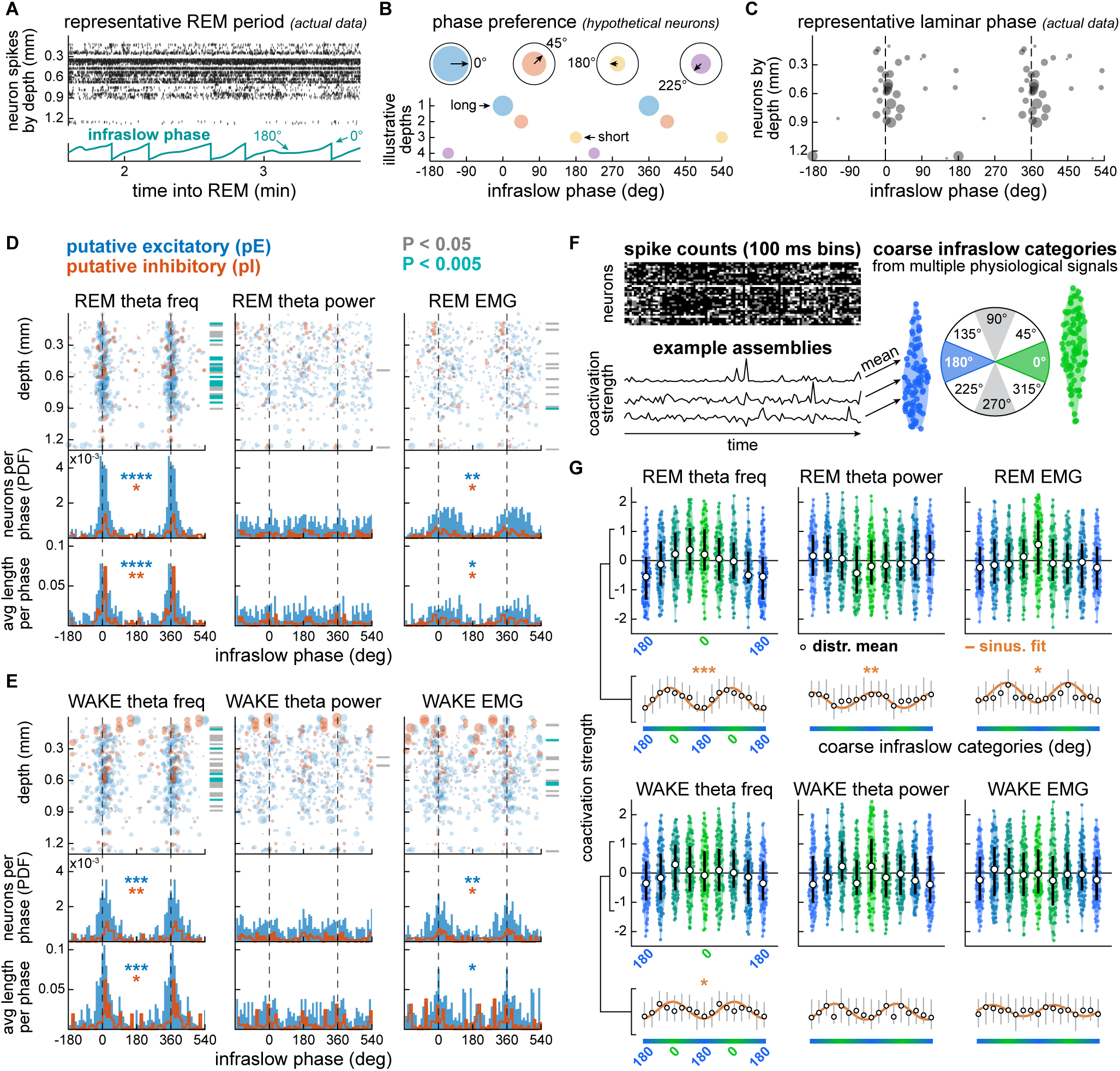
Laminar firing and spike coordination also accompany the infraslow microstructure of mouse REM. **(A)** Representative raster plot showing neuronal units along silicon probe depths (Y axis) and REM time (X axis). Black dots are spikes. We analyzed phase relationships between each unit and the Hilbert transform from infraslow fluctuations. **(B)** Hypothetical phase preference angles (X axis) and mean resultant lengths, or MRL (circle diameters) to illustrate the method by which the units in (A) were analyzed. **(C)** Scatter plot (depth vs. infraslow phase) from all REM episodes of a representative recording session, obtained using the methods in (B). **(D)** Scatter plots are similar to (C), but from all REM episodes from all home-cage sessions. Additional distinction is made between putative excitatory and inhibitory cells (pE and pI; Stark et al., 2013; Watson et al., 2016). Subplot columns show the different infraslow fluctuations. Top histograms quantify phase occupancy obtained from summing across units into 10° bins (with probability density function normalization, PDF). Bottom histograms are from averaging across MRL values. Asterisks indicate Rayleigh significance, which was robust in the phase relationship with theta frequency fluctuations (*P < 0.0042, **P < 10^−9^, ***P < 10^−21^, ****P < 10^−49^). Color-coded P values on the right of each scatter plot specify Rayleigh significance per depth. **(E)** As in (D), but from WAKE. Phase relationships with theta frequency fluctuations were weaker than in REM though still significant. Compare with Figure 4F. **(F)** Spike coordination analysis irrespective of laminar anatomy. Spikes from concatenated REM and WAKE epochs were counted in 100- ms bins per neuronal unit. Hebbian-inspired cell assemblies were then detected using an ICA-based method (Lopes-dos-Santos et al., 2013). Assembly curves were sub-epoched using the coarse infraslow categories of each given physiological fluctuation (e.g., theta frequency). Finally, sub-epoched assembly curves were averaged into coactivation strength values. Distributions of such values are shown in (G). **(G)** Each of the six subplots is from a physiological signal whose coarse infraslow categories are depicted by individual violin distributions and their mean values (white circles). The color gradient represents the infraslow continuity between 0° and 180° categories. Mean coactivation strengths seemed to fluctuate sinusoidally, especially along with REM theta frequency fluctuations, which we examined using sinusoidal fits (see below each violin graph). REM theta frequency was indeed found to drive the most significant sinusoidal patterns as quantified by Pearson’s correlations between sinusoidal fits and mean coactivation strengths (*P < 10^−3^, **P < 10^−5^, ***P < 10^−7^).

Figure 6D-E scatter plots depict the phase relationship of neuronal units with the underlying infraslow fluctuations (theta frequency, theta power, EMG). Data are from all home-cage epochs from all mice exhibiting single unit activity in their recordings (5 of 6 mice, 15 recordings, 37.4 ± 2.7 units per recording). We additionally utilized a cell classification pipeline to sort putative excitatory (pE) and inhibitory (pI) units (Stark et al., 2013; Watson et al., 2016). We quantified this by calculating probability density function (PDF) and average MRL histograms across phases (10° bins). Theta frequency fluctuations in REM were found to drive the strongest non-uniformities, according to Rayleigh tests (Figure 6D). EMG fluctuations were associated with weaker but still significant effects in both states. This was true across both pE and pI cells. No clear pattern or statistical effect emerged from theta power fluctuations (Figure 6D-E). This suggests that most cells preferentially accompany theta frequency fluctuations, regardless of the basal firing property of each cell. In addition, we determined the Rayleigh significance per linear probe channel to examine if infraslow cycles affect spiking in an anatomically heterogeneous manner, similar to our previous interrogation from LFP (Figure 5C-H). As shown by color-coded bars on the right of each scatter plot, neuronal depths indeed show varying phase relationship strengths, including with REM theta frequency fluctuations.

#### Disambiguating whisking- and theta-related spiking

One potential confound of the theta frequency results in Figure 6D-E is that the bias toward the 0° phase could be a byproduct of whisking-related spiking. In fact, our linear probe implants were made with whisker-barrel correspondence (Figure S1A-C), and we had found similar phase relationships between whisking and theta frequency (Figure 4E-F). We address this in Figure S5. We first describe that the entrainment of S1 laminar spiking to theta-filtered LFP is constant along infraslow sub-epochs, except the whisking-prone 0° sub-epochs (Figure S5C), supporting the possibility of a confound. However, according to Figure S5D, firing rates do not abruptly increase at 0° sub-epochs, but rather fluctuate constantly along the infraslow cycle of theta frequency fluctuations. Therefore, the results in Figure 6D-E most likely reflect smooth variations in firing rates rather than whisking-triggered bursts, reinforcing the intra-REM physiological continuity described in previous analyses.

#### Coordinated population activity also varies with an infraslow rhythm in mouse REM

Does coordinated spiking activity in assemblies of neurons change with REM infraslow phase? To answer this question, we used a Hebbian-inspired assembly detection method based on independent component analysis (ICA; Lopes-dos-Santos et al., 2013), thus identifying assemblies in REM and WAKE. As illustrated in Figure 6F, assembly activity was quantified in 100-ms bins, sub-epoched into infraslow categories, and averaged into coactivation strength values per category. Figure 6G is from the entire home-cage dataset and shows that assembly activity varied across infraslow categories, with different oscillating patterns emerging with respect to each physiological signal. Specifically, the pattern across infraslow categories in REM theta frequency seemed to be sinusoidal. To quantify this, we calculated correlation coefficients between each pattern and its best-fit sinusoid (see STAR methods). As shown in Figure 6G, the strongest correlation was with the oscillation of REM theta frequency, with coactivation strength peaking in the 315° category and dipping at 180°. See also Figure S6 for an alternative assembly analysis, the results of which are consistent with the findings depicted in Figure 6G.

It is worth mentioning that firing rate and coactivation strength do not coincide perfectly. Firing rates were observed to peak in the 0° infraslow category of REM theta frequency (Figures 6D and S5C-D), whereas coactivation strength peaked slightly earlier, at 315° (Figures 6G and S6). Thus it is unlikely that assembly findings were driven primarily by firing rates. We conclude that both single-unit firing rates and multi-unit assembly activity are broadly modulated by REM theta frequency, with both measures showing higher values around faster theta.

### Human REM physiology shows infraslow fluctuations that co-vary with eye movement density

Finally, we examined whether human subjects also show the continuous infraslow REM dynamics we found in mice. In Figure 7 we explore a human polysomnography (PSG) dataset from 9 healthy participants.

**Figure 7.**
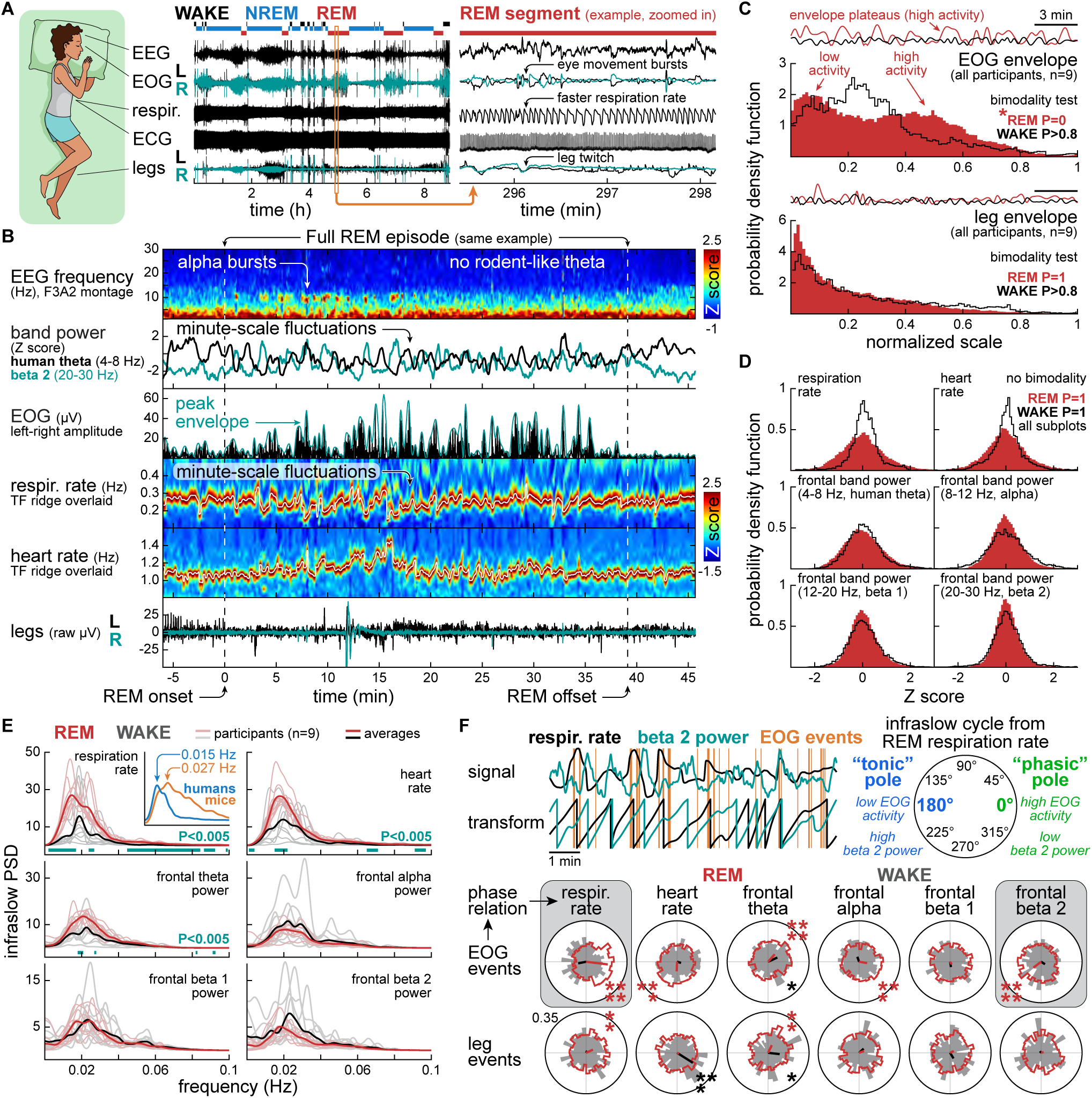
Human REM physiology shows infraslow fluctuations that co-vary with eye movement density. **(A)** Hypnogram from a full night’s sleep of a healthy participant. We explored signals from mastoid-referenced electroencephalogram (EEG), electrooculogram (EOG), respiration plethysmogram, electrocardiogram (ECG) and leg electrodes. The beige vertical rectangle delimits a REM sub-epoch, exemplifying the coincidence of eye movement bursts, faster respiration rate and leg twitches. **(B)** The same REM episode in (A) is shown in full along with NREM margins (see dashed lines indicating REM limits). Human EEG spectrograms did not show rodent-like frequency-varying theta activity, consistent with the human sleep literature (e.g., Prerau et al., 2017). Hence, theta frequency could not be used to dissect human REM microstructure with the same methods we used in mice. However, band power curves from human theta (4-8 Hz) and beta 2 (20-30 Hz) did show fluctuations suggestive of REM microstructure, similar to those in mice. Fluctuations were also observed in EOG amplitude (difference between left and right eyes), EOG envelope and time-frequency ridges from respiration and heart rate spectrograms. Leg signals occasionally showed prominent voltage events indicative of twitches (see between 10 and 15 min). See Figure S7 for additional representative epochs. **(C)** Distribution of enveloped EOG and leg signals from the entire human dataset (9 subjects). Envelopes were normalized between REM and WAKE per subject for homogenization across subjects (see representative envelopes on top of each histogram). REM but not WAKE showed significant bimodality of EOG envelopes, reflecting periods of sustained low or high oculomotor activity, consistent with the mouse saccade data (Figure 2C). Leg activity revealed no bimodality in either state when analyzed with the same enveloping method due to leg movements being more isolated. **(D)** Distribution of respiration and heart rate fluctuations, as well as frontal band power curves from the entire human dataset (9 subjects). Signals were Z scored between REM and WAKE per subject for homogenization across subjects. We found no evidence of bimodality, consistent with mouse physiology (Figure 3G-H). This further supports that REM physiology varies on a continuum, unlike EOG envelopes. Band power distributions from central and occipital montages were also unimodal (Figure S8A). **(E)** The same signals in (D) were examined for infraslow power. Rhythms slower than 0.04 Hz were ubiquitous across variables. REM showed especially strong infraslow power of respiration rate and heart rate in comparison with WAKE. The inset qualitatively compares average infraslow spectra from mouse theta frequency and human respiration rate. This shows that REM microstructure cycles at a slower pace in humans (∼67 s) than in mice (∼37 s). See Figure S8A for central and occipital data. **(F)** Phase relationships between REM sleep movements and the infraslow cycles from multiple physiological signals. Thresholded EOG events showed preference for the peaks of respiration rate cycles and troughs of beta 2 power cycles (top left). These opposite phase relationships were strongly significant, as shown by polar histograms (see highlighted by gray rectangles at the bottom). Thus, respiration rate is in-phase whereas beta 2 power is anti-phase with the “phasic–tonic” continuum (top right). This expands on findings from mice (Tort et al., 2021) and humans (Simor et al., 2020). Leg events, though more isolated than EOG events, also showed a preference for higher respiration rates, suggesting that the various motoric manifestations of human REM accompany common underlying rhythms. Rayleigh tests: *P < 0.0021, **P < 10^−4^, ***P < 10^−6^, ****P < 10^−10^. See Figure S8A for phase relationships with central and occipital band power curves.

#### Overview of the intra-REM physiology we examined in humans

The PSG signals we analyzed were: mastoid-referenced electroencephalogram (EEG), electrooculogram (EOG), respiration plethysmogram from abdomen, electrocardiogram (ECG) and legs. Figure 7A exemplifies signals from a full night’s sleep, along with the corresponding hypnogram from a sleep staging technician. In addition, Figure 7A shows a sub-epoch from the third REM episode of the same PSG session, highlighting the coincidence we often noticed between rapid eye movement bursts, periods of faster respiration rate and leg movements.

Similar to mice, we delimited human REM epochs at higher temporal resolution than the one used for sleep scoring (Figure 7B, see vertical dashed lines). As exemplified in the frontal EEG spectrogram of Figure 7B, we did not observe frequency-varying theta activity between REM boundaries, unlike mice. Rather, alpha bursts (see red blobs around 10 Hz, as reviewed by Prerau et al., 2017) and/or isolated blobs within the beta 2 range (20-30 Hz; Figure S7A, top left spectrogram) were observed in a few REM episodes of certain participants. However, intra-REM EEG patterns became more evident when examining band power curves, especially from human theta (4-8 Hz) and beta 2 bands. As illustrated in Figure 7B, human theta and beta 2 power during REM were observed to fluctuate on a minutes timescale, and roughly anti-phase relative to each other. Rapid eye movements, here measured as the absolute difference between left and right EOG and its envelope, were expectedly prominent during REM (Figure 7B). We also observed that respiration and heart rates fluctuate more strongly in REM, as evidenced by minute-scale undulating patterns in the spectrograms and their time-frequency ridges (Figure 7B). Conspicuous leg movement events, such as the one between 10 and 15 min into the representative REM episode (Figure 7B), were sparse. Figure S7 further illustrates intra-REM fluctuations in humans.

Therefore, multiple fluctuating signals suggestive of human REM microstructure were found other than frequency-varying theta activity. We next examined such fluctuations in light of our mouse analyses.

#### All human body signals show unimodal distributions, except EOG

Our first quantitative analysis of human PSG is shown in Figure 7C. EOG envelopes were concatenated across episodes per participant, both from REM and WAKE epochs (WAKE was used as control, as in mouse analyses; see Table S2 for number and duration of epochs). We then normalized the two states together per participant, thus homogenizing the data for bimodality analysis. As shown in Figure 7C (top graph), we observed two modes in the REM distribution: low and high EOG activity, the latter reflecting envelope plateaus sustained by bursts of eye movements. No such bimodality was found in WAKE. These results are consistent with our mouse data on REM eye movements (Figure 2C). Leg movements (Figure 7C, bottom) were analyzed with the same methods, except that signals from left and right legs were averaged into one combined signal prior to enveloping. No bimodality was found in REM or WAKE as leg movements were sparse, without forming plateaus of sustained activity (Figure 7C, bottom).

Figure 7D describes all other physiological signals investigated here: respiration rate, heart rate and EEG power in the human theta (4-8 Hz), alpha (8-12 Hz), beta 1 (12-20 Hz) and beta 2 bands (20-30 Hz). They were analyzed with the same Z scoring methods we used in mice. Once binned into histograms, these signals showed unimodal distributions in both states (Figure 7D). Band power distributions in Figure 7D are from frontal EEG F3A2 and F4A1 montages that were averaged together to simplify our neuroanatomical scope. The same approach was taken to analyze band power fluctuations from two additional neuroanatomical areas: central (averaged C3A2 and C4A1) and occipital (averaged O1A2 and O2A1), as shown in Figure S8A. We again observed no evidence of bimodality across distribution histograms, suggesting that the unimodality of band power curves in human REM is not neuroanatomically specific.

Therefore, among several physiological signals from human REM, only enveloped EOG suggested a bimodal REM microstructure. We observed the same contradiction between bimodal facial movements and unimodal physiology in mice (Figures 2-3).

#### Human REM physiology fluctuates at infraslow frequencies slower than mice

These same physiological signals were examined for infraslow periodicity in Figure 7E. Compared to WAKE, REM showed significantly stronger infraslow power in fluctuations of respiration rate (interaction: F_1,16_ = 18.65, P = 5.29*10^−4^), heart rate (interaction: F_1,16_ = 9.82, P = 0.006) and frontal theta band (interaction: F_1,16_ = 8.50, P = 0.010) based on two-way ANOVA with repeated measures. The inset on the top left qualitatively compares average infraslow power curves from humans and mice (X axis limits: 0-0.1 Hz, Y axis: normalized PSD). These averages were taken from human respiration rate (see main graph where the inset is) and mouse theta frequency (see Figure 3G), since these were the two intra-REM measures with the strongest infraslow PSD in this study. Thus, human and mouse REM are generally structured in cycles of 0.015 Hz (∼67 s) and 0.027 Hz (∼37 s), respectively.

#### Infraslow fluctuations in human REM are at various phases with respect to EOG bursts

Given the diversity of physiological signals fluctuating within the same infraslow frequency range, we investigated how well these signals align with the density of rapid eye movements, which is the hallmark of phasic/tonic dichotomy in human REM. As illustrated in Figure 7F (top left), infraslow fluctuations were Hilbert-transformed to extract phase at each time sample. Then, phases of EOG events were determined and plotted into polar histograms (Figure 7F, bottom; similar to Figure 4D-F). According to Rayleigh statistics, the strongest phase relationship we identified was between EOG events and REM respiration rate phase (Figure 7F, highlighted on the left). The mean phase of that relationship was at approximately zero degrees, meaning that increased EOG activity was roughly simultaneous with high respiration rate in REM. EOG activity in REM exhibited its second strongest phase relationship with fluctuations in frontal beta 2 power (Figure 7F, highlighted on the right). However, in this case the phase preference was closer to 180°, meaning that EOG activity was low when beta 2 power was high.

Because periods rich in EOG activity are traditionally recognized as phasic in human REM studies, we interpreted the zero-degree pole of respiration rate cycles as the “phasic pole” (Figure 7F, top right). So, the “phasic pole” is characterized by higher respiration rate, higher EOG activity and lower beta 2 power, with the opposite being true for the “tonic pole”. A comparable pattern emerged from our mouse analysis on theta frequency and facial movements (Figure 4D-F and Figure 5A-B). These findings concur with prior studies showing increased beta 2 power during tonic REM in humans (Simor et al., 2020).

While less prominent, significant phase relationships in REM were also found between EOG events and heart rate, as well as frontal theta and alpha power (Figure 7F, upper row of histograms). However, these relationships were not uniformly aligned to either phasic or tonic poles. Thus, different infraslow signals in REM may be at various phases in relation to rapid eye movement density.

Leg movements were also examined for phase relationships with infraslow fluctuations. As shown in Figure 7F (bottom left histogram), leg movements were more likely at higher respiration rates in REM, reflecting the same phase preference angle we had observed in the EOG-respiration relationship. Leg and EOG events also showed a similar phase relationship with frontal theta power fluctuations in REM (Figure 7F). Otherwise, REM leg movements showed statistically uniform circular distributions, which we interpret to be at least partially due to the low number of thresholded leg events. We can at least conclude that leg twitches are more likely during periods of higher respiration rate in REM, as exemplified in Figure 7A. This corroborates the phasic-tonic cycle explained above, although leg movements may be less reliable in determining the infraslow structure of human REM.

Figure S8A depicts phase relationships between EOG events and frontal, central and occipital band power fluctuations. Phase preference patterns of REM EOG with central, but not occipital, band power curves corroborated the phasic-tonic rule derived from the frontal data (Figure S8A polar histograms). This suggests that band power fluctuations from different neuroanatomical areas are not necessarily synchronized with the main REM microstructure. Furthermore, we found no parallel between REM and WAKE phase relationships, as most circular distributions in WAKE were statistically uniform, except for a few possibly spurious phase relationships in WAKE that never resembled those of REM (see all polar histograms of Figure 7F and Figure S8A). This reinforces the uniqueness of REM regarding these measures. See Figure S8B for supplemental comparisons between REM and NREM.

We conclude that there is an infraslow continuum underlying the phasic-tonic cycling in human REM, similar to mice. This phasic-tonic continuum is best synchronized with respiration rate (in phase) and beta 2 power (anti phase).

## Discussion

We demonstrate that the classically defined phasic and tonic microstates of REM sleep do not alternate bimodally at the level of physiological signals, but actually exist on a continuum that fluctuates at infraslow frequencies (0.01-0.06 Hz). We found that REM bimodality was confined to measures of facial and oculomotor activity accompanying the same infraslow rhythm, suggesting that bimodal facial movements are external expressions of a continuously varying physiology (see Figure 8 for a summary of results). Future work studying REM from the perspective of this continuous fluctuation may provide new biomarkers of sleep health, and its variation across species, developmental stages and brain disorders.

**Figure 8.**
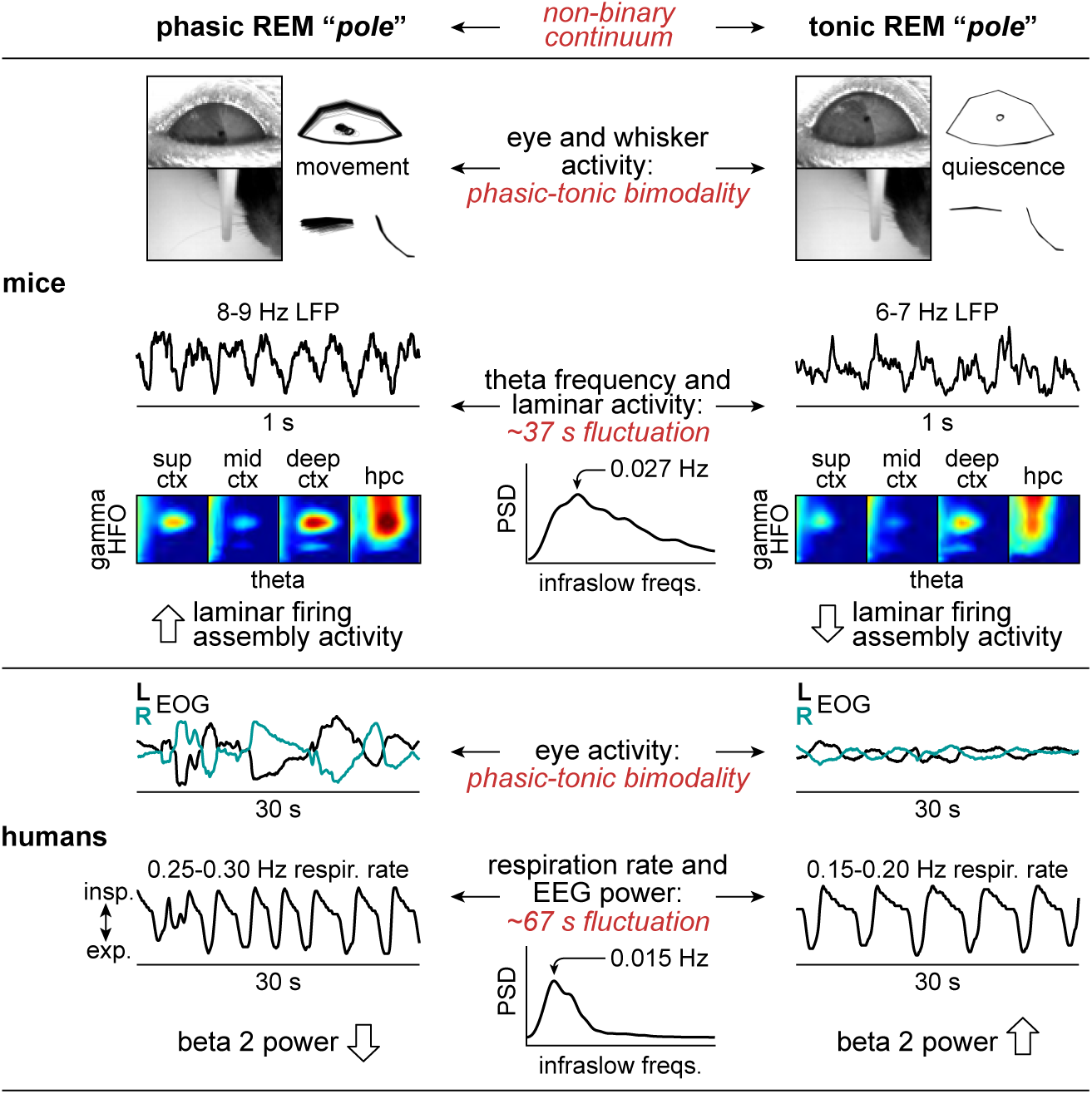
Summary of findings.

### Implications for the study of REM microstructure

The predominantly-used sleep staging method involves the division of human EEG into epochs of typically 30 s, which are then classified into the major states of vigilance, i.e., WAKE, NREM stages and REM (Prerau et al., 2017). The same methods are used to classify the 30 s periods of human REM into two microstates: phasic (high motor activity) and tonic (low or absent motor activity), with the goal of comparing them in relation to dreaming, susceptibility to awakening and physiological patterns (Ermis et al., 2010; Koroma et al., 2020; Sallinen et al., 1996; Simor et al., 2018, 2019; Takahara et al., 2002; Wehrle et al., 2007).

Rodent research conventionally adopted the same phasic-tonic dichotomy to study fine electrophysiological aspects of these supposed REM microstates (Brankačk et al., 2012; Mizuseki et al., 2011; Montgomery et al., 2008). Here we instead examined rodent REM without binary categorization. This resulted in rodent findings that may have led the way to a new understanding of REM sleep structure: that theta oscillations - a marker of rodent REM - have their peak frequency modulated at an infraslow frequency, which in turn correlates with changes in numerous other biological functions ranging from facial movement to neuronal spiking. This builds upon older studies that recognized theta frequency variations in other mammalian models, but did not quantify the periodicity of such variations (Leung, 1984; Sakai et al., 1973; Sano et al., 1973).

We then found a similar REM structure in humans, validating our mouse findings. In humans, however, this intra-REM structure was correlated with physiological measures other than theta frequency, especially respiration rate. Interestingly, respiration rate was recently reported to correlate with the frequency of brain theta oscillations in mouse REM (Tort et al., 2021). This, along with our findings, has two implications. First, we speculate that REM structure modulates - or is modulated by - several physiological fluctuations across brain and body, possibly orchestrated by common rhythm generators (e.g., respiration-related brainstem nuclei), as also discussed by Tort et al. (2021). Second, we suggest that respiration rate could be a unifying measure of human and rodent intra-REM rhythms, as opposed to hippocampal theta oscillations, which are conspicuous in rodent LFP but not clearly detectable in human surface EEG (Jacobs, 2014; Montgomery et al., 2008; Prerau et al., 2017). Thus, future studies could use respiration rate or find other physiological signals homologous between humans and rodents to better explore intra-REM rhythms, their neurochemical underpinnings, and their peripheral motor manifestations, all of which with higher temporal resolution and inter-species validation.

Another practical implication for REM microstructure derives from one of our methodological contributions: a data-driven approach to REM sub-state classification using infraslow phase coarsening. This is an alternative to imposing a binary classification of REM microstates using criteria that are often inconsistent across studies (e.g., Burgess et al., 2008; De Carli et al., 2016; Koroma et al., 2020; Mizuseki et al., 2011; Simor et al., 2016; Tamaki and Sasaki, 2019). In mice, we divided each infraslow cycle of REM theta frequency fluctuations into eight categories in an automated manner. We were then able to extract sub-epochs longer than 2.5 s (barely sufficient to analyze brain oscillations relevant to mouse REM, such as theta, gamma and HFO, as well as neuronal spiking) while still respecting the non-binary nature of intra-REM fluctuations. This clarified detailed patterns of cortical activity (further discussed below), encouraging future neuroanatomical investigation of infraslow sub-epochs, including in human REM.

### REM microstructure as a manifestation of infraslow rhythms

Infraslow rhythms are biological oscillations with frequencies between 0.01 and 0.1 Hz. These oscillations are intrinsic to a number of neurological and physiological phenomena, including heart rate, EEG power, and blood oxygen level dependent (BOLD) signal in functional magnetic resonance imaging (fMRI; Lecci et al., 2017; Picchioni et al., 2011). Accordingly, infraslow rhythms are of increasing interest among neurophysiologists studying both waking behavior and sleep (review: Watson, 2018). In fact, the resting state networks found in fMRI studies are based on cross-regional correlations occurring at the same infraslow timescale (Biswal et al., 1995; Fox and Raichle, 2007). These infraslow-based resting state networks are known to be dynamically modulated in a manner correlated with brain state (Picchioni et al., 2011), and infraslow rhythms in humans during video game performance predict behavioral performance (Monto et al., 2008; Sihn and Kim, 2022). Additionally, markers of NREM sleep in rodents and humans (including thalamic noradrenaline and sleep spindles), as well as oscillatory sequences during WAKE in medial entorhinal cortex (involved in episodic memory formation), are all coordinated at infraslow periodicity (Cogno et al., 2022; Lecci et al., 2017; Osorio-Forero et al., 2021). Thus, infraslow rhythms are becoming an increasing point of focus in cognitive and sleep neuroscience, given its omnipresence across brain states, species and techniques.

Here we incorporate REM sleep into the understanding of infraslow rhythms. Two previous studies had actually noted minute-scale periods in human REM: one EOG study reporting that the density of rapid eye movements fluctuates at about 2 minute intervals (Ktonas et al., 2003), and an fMRI study describing a fluctuation in resting state network signals during REM at a frequency of 0.01-0.1 Hz (Chow et al., 2013), both of which overlapping with the infraslow frequencies of our findings. Our work, however, is the first to explicitly demonstrate the infraslow periodicity in REM through inter-species comparisons and the use of several controls, including the phase relationship between physiological fluctuations and rapid movements in mice and humans, which identified infraslow signals optimally related with “phasic” and “tonic” motricity in each species (theta frequency in mice, respiration rate in humans).

Since we implanted linear probes across neocortical layers and hippocampus in mice, we were also able to map electrophysiological measures onto both a spatial (laminar depths) and temporal continuum (REM infraslow sub-epochs). We found that theta phase optimally modulates HFO amplitude at a particular location (deep neocortex) and timeframe (“phasic REM pole”). This links two previous studies that separately analyzed the dependency of phase-amplitude coupling on the anatomical depth (Scheffer-Teixeira and Tort, 2017) or binarily-classified REM microstates (Brankačk et al., 2012). In addition, we observed that the infraslow microstructure of REM affects two measures of spiking activity: firing rates across neocortical/hippocampal depths and the tendency of spikes to fire as coactivating assemblies of neurons (Lopes-dos-Santos et al., 2013). We again found these to be higher around the here-proposed “phasic REM pole”, consistent with the notion that phasic REM creates opportunities for transregional coordination (Montgomery et al., 2008). Based on our contribution, all these processes can now be examined under the infraslow time metric, which will potentially optimize the prediction of network function variations within REM, as well as their emergent properties, such as the different stages of memory consolidation. Future studies could also formally examine the laminar anatomy of intra-REM fluctuations by assigning neuronal units to physiological layers, as done by Senzai et al. (2019).

### Involvement of within-REM rhythms in development and sleep health

What other benefits might within-REM rhythms serve? One conception of REM comes from studies on its muscular twitches, which emerge developmentally as one of the first coordinators of neural activity in the young brain. It is hypothesized that REM twitches enable sensorimotor feedback early in infancy, thus promoting activity-dependent circuit sculpting in preparation for further plasticity later in life (Blumberg et al., 2013; Dooley et al., 2020; Sokoloff et al., 2020; Tiriac et al., 2012). Thus, early-life within-REM rhythms may allow network tuning to internally versus externally driven activity in a cyclical manner, and the persistence of such cycling in adult REM may be important for continuing circuit adjustment. Further investigation on these REM cycles and their impact on both behavior and neural activity during WAKE may be amenable to modeling and experimental approaches.

Healthy phasic-tonic cycling may also play important roles for REM-related processes such as homeostatic effects on neuronal spiking (Grosmark et al., 2012; Watson et al., 2016), dreaming (Siclari et al., 2017) and consolidation of emotional memories (Vandekerckhove and Wang, 2018). In particular, over-consolidation of emotional memories, as is the case in post-traumatic stress disorder (PTSD), as well as major depression, have been proposed to involve alterations in REM sleep, in addition to alterations in sleep architecture more generally (Murkar and De Koninck, 2018; Riemann et al., 2020). Furthermore, dysfunctional phasic and tonic muscle activity in REM sleep behavior disorder (RBD) predicts neurodegenerative disorders, such as Parkinson’s disease (McCarter et al., 2014; Schenck et al., 1996; St Louis et al., 2017). Also interestingly, abnormal separation between phasic and tonic REM features, possibly indicative of dysfunctional intra-REM cycling, was found to predict the conversion to neurodegenerative disease from RBD (Nepozitek et al., 2019). Therefore, better analytical methods to characterize human REM, such as the ones proposed here, combined with accessible measures, such as respiration rate, may provide greater diagnostic or prognostic information.

## Conclusion

Our findings deepen our understanding of REM sleep and its internal heterogeneity. This is important given that REM is the least explored of the three major arousal states of WAKE, NREM and REM. The discovery here of infraslow-modulated variations in brain and body signals may provide a foundation on which to study how network functions vary within REM. Finally, previous linkages of REM microstates to disease may be more systematically studied using this quantitative biomarker of healthy REM.

## Acknowledgments

We thank colleagues from the Michigan Neuroscience Institute: Dr. Christian Burgess for inputs on eye videography and head-fixed sleep in mice; Drs. Daniel Z. Doyle and Mandy M. Lam from Dr. Kenneth Y. Kwan’s lab for guidance on immunofluorescence histology. We also thank colleagues from VA Portland Health Care System and Oregon Health and Science University: Peyton T. Wickham and Dr. Christy Reynolds from Dr. Miranda M. Lim’s group for their assistance with the human polysomnography pipeline. We thank all members of Dr. Brendon O. Watson’s lab for internal discussions, especially former lab member Mingxin Ding for starting our head fixation setup. We finally thank Dr. Eric Simon of the Research Innovation Core at the University of Michigan’s Eisenberg Family Depression Center for reviewing and editing this manuscript. This work was supported by NIH grant MH107662 and the University of Michigan Neuroscience Scholars fund to B.O.W., and the Portland Veterans Affairs Research Foundation to M.M.L.

## Author contributions

Conceptualization, B.O.W., L.S.B.J. and M.M.L; Methodology, L.S.B.J.; Software, L.S.B.J., M.S.R. and B.O.W.; Formal Analysis, L.S.B.J. and M.S.R.; Investigation, L.S.B.J.; Resources, B.O.W. and M.M.L.; Data Curation, L.S.B.J., M.S.R. and M.M.L.; Writing - Original Draft, L.S.B.J.; Writing - Review and Editing, B.O.W., M.S.R. and M.M.L.; Visualization, L.S.B.J.; Supervision, B.O.W.; Project administration, B.O.W.; Funding Acquisition, B.O.W.

## Methods

### Mice

We used six adult male mice (C57BL/6, The Jackson Laboratory) with free access to food and water, 12-hour light/dark cycle (lights on at 6 AM), and standard conditions of bedding, temperature, humidity and ventilation. Animals were 9-10 weeks old upon arrival at the vivarium and were group-housed until the implantation surgeries, performed at 11-13 weeks of age. After surgeries, the mice were single-housed for the remainder of the study. Experiments were carried out in the A. Alfred Taubman Biomedical Science Research Building at the University of Michigan Medical School in compliance with bioethical guidelines from the National Institutes of Health (protocol PRO00009818, approved by the Institutional Animal Care and Use Committee).

### Humans

De-identified polysomnography (PSG) data from human subjects were retrospectively analyzed under the Veterans Affairs Portland Health Care System (VAPORHCS) Institutional Review Board approval (MIRB #4108), a larger study which was conducted in accordance with the ethical guidelines of the Belmont Report. Participants underwent in-lab, attended PSG (details below) after referral to the VAPORHCS Sleep Clinic between 2018 and 2021 (n = 9). Deidentified data were curated, exported, cleaned and transferred to the University of Michigan Medical School under an approved Data Use Agreement between institutions. Subjects were included if they: (1) were 18-70 years of age; (2) underwent diagnostic-type PSG with greater than 6 hours of sleep recorded overnight; showed more than 25 minutes of total REM time across two or more discrete periods scored over the entire night; had an apnea-hypopnea index of less than 5 (e.g., no clinically significant sleep apnea); (5) had a periodic limb movement index of 0; (6) were otherwise healthy and did not endorse traumatic brain injury, posttraumatic stress disorder or depressed mood using Patient Health Questionnaire-9 and (7) had less than 5% of epochs containing movement or other signal artifact per visual inspection.

### Surgeries and implant validation

Each mouse underwent two surgeries one week apart. In the first surgery, we performed intrinsic optical imaging of S1 barrel cortex followed by partial construction of the chronic head cap. The second surgery aimed at imaging-guided silicon probe implantation followed by completion of the head cap.

#### Intrinsic optical imaging and head cap base

Imaging procedures were mostly based on Aronoff and Petersen (2007) and Knutsen et al. (2016), with the aid of a stereomicroscope on an articulating stand (Leica). Mice were anesthetized with isoflurane (1-1.5% v/v in oxygen; VetOne) and maintained at 37 °C on a heating pad adapted to the stereotaxic frame (Kopf). Prior to ear-bar fixation, all whiskers on the right side of the animal were cut except six: C1-2, D1-2 and E1-2, which were reserved for later stimulation. We then proceeded with standard midline incision to expose bregma and lambda. A skull area above the left barrel cortex was skim-cut using a drill, forming a square thinned-skull window. The size and position of this window were based on canonical barrel coordinates (Figure S1A; Knutsen et al., 2016) corresponding to the spared whiskers. Once ready, the skull window was maintained wet with saline and covered with a glass coverslip for reversible skull transparency.

Images of the skull window were obtained using a 12-bit grayscale camera (MV1-D1024E-G2, PhotonFocus) attached to a VZM 200i lens (Edmund Optics) angled 30° in relation to the sagittal plane to match the curvature of S1 cortex. For illumination, we used a lab-assembled high-power LED setup (Roithner LaserTechnik) with manual switch between red light (630 nm) for functional imaging of the brain surface and green light (525 nm) to collect an anatomical reference image highlighting the blood vessels on the same surface. Whisker stimuli consisted of 10-Hz sinusoidal vibrations on the rostral–caudal axis.

Stimulation was performed one whisker pair at a time (C1-2, D1-2, E1-2) by inserting them into a glass micropipette attached to a piezoelectric actuator (Piezo.com) controlled by a biphasic stimulus isolator. This hardware was mounted on a custom setup of platforms and stands (ThorLabs). Camera and piezoelectric actuator were jointly triggered by a Bpod State Machine (Sanworks) via custom implementation of the Bpod code repository.

Red-illuminated frames were collected continuously at 4 Hz throughout a sequence of 40 trials, each trial with 4-s baseline, 4-s whisker stimulation and 4-s post-stimulation period. Using Matlab (Mathworks), individual stimulation frames (stim) and baseline frames (no-stim) were normalized against the average baseline image of that trial (Knutsen et al., 2016). Normalized frames were then averaged across trials, and across stim or no-stim, respectively producing images with and without brain surface signals (Figure S1A). By repeating this 40-trial experiment one whisker pair at a time, we obtained their respective stim/no-stim images. Brain surface signals were smoothed, thresholded (pixels beyond 2 standard deviations were classified as signal) and merged onto the blood vessel snapshots (Figure S1A), creating mouse-specific stereotaxic maps for later use.

Once brain imaging was finished, the glass coverslip used for skull transparency was removed. Four microscrews were placed into the bone around the thinned-skull window. A fifth burr hole over the contralateral cerebellum received a male-female connector pin for future electrical grounding (22 AWG; male portion touching the cerebellum). Then a 3D-printed RIVETS head fixation ring (Figure 1A; Osborne and Dudman, 2014) was placed on the skull around the microscrews and ground pin. Ring, microscrews and ground pin were finally cemented together (C&B-Metabond adhesive cement, Parker). The thinned-skull window was left exposed without cementation, temporarily protected by a drop of silicone sealant (Kwik-Cast). Analgesic anti-inflammatory treatment (Carprofen, 5 mg/kg, SC) was administered 1 h prior to concluding the surgery and a one-week post-surgical recovery was allowed.

#### Imaging-guided implantation

For the second surgery, the setup was converted from imaging to anesthetized electrophysiology. This system included a micromanipulator (Sensapex) holding the combination of silicon probe (one linear shank, 64 electrode sites on edge of shank, 20-μm spacing between sites; NeuroNexus), amplifier/digitizer headstage, and recording cable leading to a multichannel interface, all within Faraday protection (electrophysiology is described later in this text). Prior to the second surgery, intrinsic imaging maps were inspected to identify the strongest neocortical response per mouse (Figure S1A). The selected map was kept on display on a computer monitor to guide the implantation.

Mice were again anesthetized and fixed by the ear bars. The stereomicroscope was positioned for visualization of the thinned-skull window after removing its silicone sealant protection. Using the micromanipulator, we pointed the silicon probe at the area of the skull window corresponding to the neocortical response at a 30° angle in relation to the sagittal plane. The resulting antero-posterior and meso-lateral coordinates were locked from this point on. The probe was then moved away from the mouse for safe removal of the thinned skull. Ground wires from the probe connector package were plugged into the cerebellum pin and the recording was turned on. The probe was lowered back toward the brain until touching the dura mater, as indicated by a cross-channel voltage deflection, at which point the dorsal-ventral coordinate was calibrated to zero. We kept lowering the probe against the brain, attempting to perforate the dura mater with the probe itself. In case of resistance and risk of breakage of the probe, we retracted the probe and carefully pierced the dura mater using an insulin needle before resuming the implantation.

The final dorsal-ventral trajectory of the probe was made at constant speed (1 mm/min) with real-time monitoring using an open-source electrophysiology viewer (Neuroscope). The viewer was pre-configured with a channel map corresponding to the order of electrode sites on the probe. This allowed us to better visualize the transitions from electrical noise to brain signal on a channel-by-channel basis as electrode sites gradually entered the brain (Figure S1B, left). The dorsal-ventral trajectory was interrupted as soon as the most superficial channel started displaying brain signals.

Once the implantation was finished, the probe shank was entirely covered with heated vaseline. We then built up the head cap around the vaseline, connector package and ground wiring using regular acrylic resin. The surgery was concluded by disconnecting the headstage from the probe and releasing the mouse. Analgesic anti-inflammatory treatment (Carprofen, 5 mg/kg, SC) was again administered 1 h prior to concluding the surgery and an additional week was allowed for post-surgical recovery.

#### Somatotopic verification

One assumption from our surgeries was that we had achieved whisker-barrel somatotopy given the S1 imaging-based implantation we used. To verify this electrophysiologically, we performed an awake whisker stimulation experiment under head fixation (Figure S1B, center). This experiment was made after surgical recovery prior to the main sleep recordings (the head-fixation hardware is described later).

Briefly, animals were head-fixed for manipulation of the spared whisker pairs: C1-2, D1-2 or E1-2 (right side of animal). A randomly selected whisker pair was reversibly glued onto a custom 3D-printed pole, horizontally movable by a linear actuator (NA series, Zaber). Using voltage triggers from Bpod State Machine (Sanworks), we stimulated the whisker pair by moving it 4 mm caudally or rostrally in random order (stimulus duration: 200 ms, delivered every 5 s) for 15 min (180 trials total). The procedure was repeated for the remaining whisker pairs, always with timestamping of stimulus triggers. We performed this experiment in a single session, once per mouse.

After recordings, electrophysiological epochs from all 64 silicon probe channels were extracted around whisker stimuli (100 ms before, 300 ms after stimulus onset), resulting in 2D matrices with channels in rows and digital samples in columns, one matrix per trial. Using Matlab, matrices were averaged and analyzed as CSD (Chen et al., 2011) by calculating the second derivative across channels (built-in *gradient* function, used twice) followed by spatial smoothing (10-channel window) and temporal smoothing (20-ms window; Figure S1B, center). Because the three whisker pairs had been stimulated one at a time, we were able to separately analyze their CSD response strength as measured by the mean absolute response across both dimensions (channels and time). The whisker pair yielding maximal response was spared for the remainder of the study, assuring extra somatotopic targeting. All other whiskers were trimmed weekly under isoflurane anesthesia. Ipsilateral whiskers were not manipulated throughout the study.

#### Cortico-hippocampal depth profile

In addition to implanting across the S1 barrel cortex at a 30° angle, we assumed that the probe had reached into an underlying subcortical area since the length of the electrode array (1.28 mm) was known a priori to exceed the thickness of S1 (∼0.96 mm; Paxinos and Franklin, 2019). That subcortical area was supposedly the lateral portion of the dorsal hippocampus since it is diagonally aligned with the medial barrel cortex, especially around -1.5 mm from bregma (Paxinos and Franklin, 2019). To verify this depth profile, we examined theta ratio and high-frequency oscillation (HFO) band power (Figure S1B, right). We performed this particular analysis on home-cage recordings. For laminar theta ratio, we divided theta power (5-10 Hz) by the power of a wider theta-surrounding band (2-20 Hz) per channel. For laminar HFO, we simply obtained HFO power (100-160 Hz) per channel (Figure S1B, right).

#### Histological verification

After experiments, each mouse was deeply anesthetized with isoflurane (5% v/v in oxygen) and underwent standard brain perfusion techniques, including diaphragm incision to expose the heart and right ventricle cutting. We injected 5 mL of phosphate-buffered saline (PBS) and then 5 mL of 4% paraformaldehyde (PFA) into the left ventricle. After decapitation, the brain was removed from the skull and stored in 4% PFA under 4 °C for at least 24 h. Using a vibratome, each brain was sectioned at 60-µm thickness around the antero-posterior level containing S1 barrel cortex (Paxinos and Franklin, 2019). Sections were placed in PBS-filled wells of a tissue culture plate and examined for silicon probe tracts using a stereo microscope. Four tract-containing sections were selected for tissue processing.

Selected sections were orbitally shaken for 1 h in a culture plate with detergent-containing blocking solution (5% donkey serum, 1% BSA, 0.1% glycine, 0.1% lysine and 0.3% Triton X-100 in 1x PBS) in order to permeabilize membranes and reduce nonspecific antibody binding. Sections were incubated overnight with primary antibodies anti-NeuN (chicken-raised, RBFOX3, cat. number - ABN91, Millipore-Sigma) and anti-Netrin-G1a (goat-raised, cat. number - AF1166, R&D Systems) in a cold room and then incubated for 2 h with fluorophore-containing secondary antibodies at room temperature (Donkey anti-Chicken IgY (H+L), Alexa Fluor 488 Affinipure - Jackson ImmunoResearch Labs 703-545-155; Donkey anti-Goat IgG (H+L), Alexa Fluor Cy3 AffiniPure - Jackson ImmunoResearch Labs 705-165-147), both under orbital shaking. DAPI was added to the secondary antibody step, enabling triple staining. Sections were placed on glass slides and covered with an immunofluorescence mounting medium (Vectashield). Specimens were observed using an immunofluorescence system composed of a dissecting scope (Olympus SZX16), fluorescent source (Olympus U-HGLGPS), camera (Q-imaging Retiga 6000) and imaging software (QCapture Pro 7). Images were acquired at 8-bit depth, 2×2 binning and variable exposure depending on the fluorescence wavelength. Finally, images were colored (Figure S1C) using an open-source software (Fiji).

### Extracellular electrophysiology in mice

Brain activity was amplified (1000x) and digitized (20 kHz) by a 64-channel headstage and transmitted through an SPI cable into a multichannel interface with USB connectivity (RHD2000 system, Intan). All electrophysiological data were processed and analyzed in Matlab using custom implementation of built-in functions, public code repositories and online. This study also resulted in its own code repository (unpublished as of the posting date of this preprint). The wide-band signal was preprocessed as local field potentials, LFP (450-Hz low-pass filtering, sampled at 1.25 kHz), electromyogram (EMG) from intracranial signals (Schomburg, 2014; 300-600 Hz filtering followed by pairwise channel correlations, here sampled at 1.25 kHz) and spiking activity (500-Hz high-pass filtering, sampled at 20 kHz).

### Home-cage and head-fixed hardware for mice

Home-cage sleep sessions consisted simply of allowing the mice to move and sleep freely in their own cages while plugged into the recording cable (Figure 1A). A tall cage lid without the filter top was used instead of the regular lid, allowing passage of the cable while preventing animals from escaping. For Faraday protection, the cage was reversibly wrapped in aluminum foil or copper mesh, which was wire-connected to the ground via the multichannel electrophysiological interface. No changes were made to the bedding or environmental enrichment of the cage.

Head-fixed sleep was recorded from an electrophysiology and videography setup centralized by a pair of head-fixation forks adapted from the RIVETS system (Figure 1A-B; Osborne and Dudman, 2014). While head-fixed, mice stayed on a custom 3D-printed platform with walls: the “sleep box” (dimensions: 11 × 6 × 2 cm; designed in SolidWorks). This sleep box was 3D-printed using white resin (FormLabs), providing a white background for video recording of the right whiskers using a camera positioned above the mouse (13-cm distance between lens and mouse). For eye videography, we used a laterally positioned camera (7-cm distance between lens and mouse) framing the left eye through an aperture in the wall of the sleep box (head-fixed videography is described in detail later). For illumination, we used two infrared LED boards (inexpensive LED used in security cameras). These components were assembled using a custom setup of platforms and stands (ThorLabs). Grounding was achieved by connecting sources of electrical noise to a surrounding metal mesh, which in turn was wire-connected to the ground input of the multichannel electrophysiological interface (Intan).

### Mouse sleep scoring and REM epoch refinement

Our main recordings started regularly at 6:30-7:30 AM, i.e., at the beginning of the light period, and lasted 387 ± 10 min (mean ± standard error). Sleep scoring was made in 1-s bins based on published spectrogram-based procedures (Grosmark et al., 2012; Watson et al., 2016) using a community-maintained algorithm (*SleepScoreMaster*, buzcode repository). Three variables were used for scoring: power spectral slope (PSS; i.e., value of the negative slope formed by power spectra across log-spaced 4-100 Hz frequencies), theta ratio and normalized EMG. The 1-s bins typically formed three clusters, each one representing a brain state: NREM (low PSS), REM (high theta ratio, low EMG) and WAKE (all other times). PSS and theta ratio were respectively obtained from one neocortical and one hippocampal channel per session. The assigned neocortical channel was the one showing strongest sensitivity between WAKE and NREM spectral states as indicated by the bimodality of PSS distributions. The assigned hippocampal channel was the one with the strongest theta ratio.

The next step was to manually curate the sleep scores. We used a community-maintained GUI tool (*TheStateEditor*, buzcode repository) comparing WAKE, NREM and REM labels against an interactive spectrogram (1-40 Hz, linearly spaced frequencies) accompanying theta ratio and normalized EMG curves (Figure 1C). This curation involved minor corrections of state boundaries and, most importantly, the extension of REM episodes in both directions, creating 30–60-s margins for later refinement.

Margins around REM were manually trimmed using a second GUI developed during this study. The GUI displayed wavelet spectrograms from REM episodes one episode at a time (Figure 1D). Each REM spectrogram was generated at higher temporal resolution (125 Hz instead of 1 Hz) with linearly spaced frequencies around the theta band (0-15 Hz, 0.05-Hz bin size). Each frequency bin was assigned a Morlet wavelet kernel to convolve the LFP epoch (*MorletWavelet* and *FConv* functions, buzcode repository; based on Cohen, 2014). The absolute of the resulting matrix was smoothed along frequency (1-Hz window) and time (5-s window) and Z scored along frequency.

By interacting with these spectrograms via GUI, we manually timestamped the onset and offset of each REM episode. REM onsets consisted of relatively sharp transitions from slow-wave and spindle activity (1-15 Hz) to highly concentrated theta activity (5-10 Hz), replicating the literature (e.g., Bandarabadi et al., 2020; Grosmark et al., 2012). REM offsets, in turn, consisted of transitions from strong theta oscillations back to slow-wave activity (Figure 1D). Once finished looping through REM episodes, we obtained lists of refined REM epoching timestamps, one list per recording session. These epoching timestamps were used in the remainder of the study.

Based on the list of REM timestamps of a given recording session, we produced an equivalent list for WAKE epoching. Thus, for each REM episode we delimited a paired WAKE epoch with the exact same duration so that REM and WAKE could be statistically compared in an own-control design (Figures 2 and 3). WAKE was considered an important control given its incidence of theta and motor activity, comparable with REM. Control WAKE epochs were extracted from the initial hour or so of recording, during which the theta ratio was typically stronger and animals were more active (Korotkova et al., 2018). We avoided periods contaminated by movement artifacts when creating these WAKE epochs to maximize comparability with REM epochs, which were mostly artifact-free.

### Head-fixed videography in mice

#### Acquisition

The left eye was filmed using a 10-bit grayscale camera (FL3-U3-13S2M, Flir) coupled with a VZM 200i lens (Edmund Optics). Illumination consisted of infrared reflectance using a regular LED board, without the trans-skull back illumination method used in previous studies (e.g., Sattler and Wehr, 2021; Yüzgeç et al., 2018). Acquisition was mediated by a 3.0 USB card (U3-PCIE2-2P01X), with frame rate (20 frames/s) and frame size (640 × 512 pixels) configured using the FlyCapture software (Flir). Videos were written into AVI files (M-JPEG codec) using the same software. The camera was additionally used as a source of digital timestamps (1-ms duration), one per frame. These timestamps were streamed into the multichannel electrophysiology interface (RHD2000 system, Intan) for time-locked analysis along with all other data at a 20 kHz sampling rate.

Right whiskers were filmed against an infrared-illuminated white background using an 8-bit grayscale camera (EoSens CL, Mikrotron) coupled with a 1:1.8/50 mm lens (CF50HA-1, Fujinon). Image acquisition was mediated by a frame grabber (Xtium-CL MX4, Teledyne Dalsa), with frame rate (100 frames/s) and frame size (512 × 320 pixels) configured using the MC ControlTool software (Mikrotron). Videos were recorded into mp4 files (H264 codec) using the StreamPix software (NorPix). The frame grabber was used to generate the digital timestamps (1-ms duration) representing frames. These timestamps were also sampled at 20 kHz via the multichannel interface (Intan).

#### Video epoching

Eye and whiskers were filmed continuously to capture the spontaneously occurring REM episodes. Once the electrophysiological sleep scoring was finished, we were able to extract REM-containing video segments (or their WAKE counterparts), thus avoiding the expense of analyzing the entire videos. To extract these video segments, we delimited the camera timestamps within each REM or WAKE epoch (at the electrophysiological sampling rate) and indexed the corresponding frames in the original video file (built-in *VideoReader* and *read* functions). Indexed frames were then written into the new epoched videos (*VideoWriter* function).

#### Tracking of body parts

Facial movements were tracked using DeepLabCut, or DLC (Nath et al., 2019). To train the network that would later be used to analyze eye and pupil (20 frames/s), we provided DLC with manually selected frames representing a diversity of eye video scenarios (Figure S2A). They varied from easy, unobstructed imaging of eyelids (whether blinking or not) and pupil (whether moving or not, whether dilated or not) to challenging situations, such as pupil hidden behind the eyelids or a small amount of eye secretion, in which case we would miss pupil movements but still be able to record eyelid movements. Once representative frames were selected, we placed eight labels on the eyelids and eight labels around the pupil, forming two octagons (Figure 2A). Pupil labeling was skipped in frames with closed eyelids. We then trained the network overnight.

A similar process was carried out when preparing the whisker tracking network. Video situations varied from idle whiskers with good focus to moving whiskers with motion blur (Figure S2B). Motion blur was due to the frame rate we used in this study (100 frames/s), i.e., lower than the high-speed videography employed in whisker-oriented studies (>200 frames/s; e.g., Cheung et al., 2019). Nevertheless, the frame rate we used was deemed a good tradeoff between video file size - important, given our several hour-long recordings - and the temporal resolution needed to capture REM twitches (Video S1). Once representative frames were selected, we placed four labels on each whisker (labeling was skipped in frames with whiskers out of focus), and six labels on the nose and mystacial pad (Figure 2A). Nose and mystacial pad labels were used only for visual reference and were not analyzed.

Once trained, DLC networks were used to track each video epoch. Labeling accuracy was evaluated one video epoch at a time using quality inspection tools provided by DLC: plots showing cumulative body part trajectories and demonstration videos showing label positioning in real time (Nath et al., 2019). Approved video epochs had their data submitted to a second layer of analysis (see below). Problematic videos, e.g., with excessive eye secretion, were excluded (Table S1).

#### Eye and whisker activity

DLC labels consisted of X and Y coordinate pairs across the pixel grid of each video frame. We converted these pixel coordinates to millimeters using calibration snapshots we had taken from a ruler prior to analysis. Millimeter coordinates were then analyzed as polygons around eye/pupil and polylines on whiskers (Figure 2A, Video S1).

These geometric shapes allowed multiple measures across the time axis (Figure 2B), explained as follows. Eyelid opening: roughly vertical distance (hypotenuse) between the dorsal-most and ventral-most eyelid vertices, followed by moving average (5-s window) to attenuate spike-like events corresponding to blinking. Pupil diameter: roughly horizontal distance (hypotenuse) between the anterior-most and posterior-most pupil vertices. Saccading: speed of the dorsal-most pupil vertex, calculated as its shift between adjacent frames (hypotenuse) divided by the time between frames. Here we present speed in millimeters per decisecond (mm/ds) since individual saccades were observed to be on the scale of tenths of seconds (Video S1). Blinking: the raw eyelid opening curve minus its own moving average (5-s window), revealing blinks as spiky events. Whisking: same speed calculations used for saccading but from the multiple vertices on the whiskers. Speed curves from the multiple whisker vertices were averaged into a single curve representing whisking activity, also in mm/ds. Because speed curves from individual whisker vertices were mostly redundant, their averaging was useful to smooth out irregularities from erratic DLC labels in a minority of frames with motion blur (Figure S2B). This redundancy was also useful to homogenize whisker speed curves across videos with single (Figure 2A) and double whisker tracking (Figure S2B).

#### Distributions of enveloped rapid movements

Saccading, blinking and whisking curves suggested a temporal structure, with alternation between movement bursts and quiescence (Figure 2B). We captured this alternation using peak envelopes (2-s window; built-in *envelope* function), which were then examined for their distributions across normalized scales (number of bins per histogram: 100; Figure 2C). Normalization was achieved by jointly rescaling duration-matched REM-WAKE epochs from 0–1. The same normalization and histogram binning was performed on EMG data (125 Hz), allowing us to compare myoclonic activity and nuchal tone.

### Mouse theta activity

#### Paired theta spectrograms

For each REM and WAKE epoch, we generated a wavelet spectrogram using the same sampling rate (125 Hz) and methods explained above. This time, however, spectrograms were focused on theta (5-10 Hz) and jointly Z scored within each REM-WAKE pair along the frequency dimension (Figure 3C).

#### Theta frequency vs. theta power

Wavelet spectrograms yielded two separate measures of theta activity: frequency and power. They were respectively calculated using maximum-energy time-frequency ridges (*tfridge* function) and frequency bin averaging.

Ridges consisted of frequency bin values along the spectrogram time axis. Ten ridges were extracted from each spectrogram, each ridge surrounded by one frequency bin. Thus, once a ridge and the adjacent frequency bins were extracted in iteration *i*, they would be ignored in iteration *i*+1 for extraction of the next ridge. Ridges were then smoothed (5-s window; Figure 3D). This procedure allowed us to compare REM and WAKE in terms of the contiguity among multiple ridges. To translate this contiguity into a single curve per spectrogram, ridges were divided by their mean variance within that spectrogram and averaged (Figure 3E, top and center). This resulted in new curves negatively weighted by ridge variance, i.e., the lower the ridge variance, the higher the curve values (in arbitrary units).

Theta power fluctuations were simply obtained by averaging across the frequency bins of each spectrogram prior to Z scoring them. All ridge and power curves were mean-subtracted, one epoch at a time. Curves were then Z scored within their respective REM-WAKE pairs, generating zero-centered fluctuating patterns with comparable amplitudes between theta frequency and theta power (Figure 3E).

#### Infraslow spectrum and distribution

The aforementioned fluctuations were observed to occur in cycles of several seconds, meaning that short epochs (e.g., <1 min) usually contained no more than 2-3 complete cycles. Therefore, we decided to concatenate across epochs, creating minute- to hour-long oscillatory curves (Figure 3F). With these concatenated curves, we constructed sufficient oscillatory data for infraslow (<0.15 Hz) spectral analysis. For such spectral analysis, we calculated Welch’s power spectral density, PSD (built-in *pwelch* function, 3-min Hamming window, 50% overlap) at infraslow frequency bins (0–0.15-Hz band, 0.001-Hz bin size). We obtained one pair of REM and WAKE PSD curves per mouse, i.e., each PSD was calculated from all concatenated REM or WAKE epochs of that given mouse (Figure 3G-H). The same concatenated epochs were submitted to distribution analysis using histograms (100 bins) across Z score scales (Figure 3G-H).

### Combined video and electrophysiology in mice

#### Correlations and clustering

We took advantage of the head-fixed dataset to co-examine theta frequency ridge, theta power, EMG and enveloped saccading, blinking and whisking. Our first approach was to explore Pearson’s linear correlations among these six variables. First, we standardized the different sampling rates of these variables (electrophysiology: 125 Hz; eye video: 20 Hz; whisker video: 100 Hz) to a common binning (2-s bin size). Bins were then concatenated across all epochs from all mice (after excluding problematic videos; Table S1) and examined for pairwise correlation coefficients. Separate correlation matrices were created from REM and WAKE (Figure 4A).

A second analysis was performed on the 2-s bins, particularly from the three facial movement dimensions: enveloped saccading, blinking and whisking. First, we scatter-plotted these three dimensions and observed them to form two overlapping clusters per brain state (Figure 4B). Then we fitted a two-component Gaussian mixture model to REM and WAKE data separately (built-in *fitgmdist* function). Based on the mean and standard deviation of each model component, we constructed ellipsoid polyhedrons (*PLOT_GAUSSIAN_ELLIPSOID* function). These polyhedrons were then used to index the data points as belonging to either one of the model components or none of them (using built-in *delaunayn* and *tsearchn* functions). This resulted in new clusters excluding ambiguous and peripheral bins. Clusters with lower and higher mean normalized values were identified as facial quiescence and facial movement, respectively (Figure 4B). Finally, these clusters were compared within REM and WAKE in terms of electrophysiological measures using violin plots.

#### Facial movement phase relationships

In addition to the 2-s binned analysis explained above, we investigated the timing of raw (not enveloped) facial movement events in relation to the infraslow cycles. The first step was to upsample video data to 125 Hz using linear interpolation, thus matching the sampling frequency used for the electrophysiological data. Spike-like events in saccading, blinking and whisking curves were then thresholded into timestamps (Figure 4D; like Lerma and García-Austt, 1985). The threshold we used was the mean saccading, blinking or whisking value of a given epoch plus 2x the standard deviation (manually adjustable by a custom GUI on an epoch-by-epoch basis). Timestamps were taken from upward threshold crossings, resulting in one timestamp per event.

Next, theta frequency ridge, theta power and EMG were analyzed as the phase angles (0 to 2π rad) of their Hilbert transforms (built-in functions *hilbert, angle* and *mod*). Movement timestamps were used to index into these phases, thus capturing phase relationships between movement events and the transforms. Circular distributions from event-indexed phases were then normalized against their underlying transform distributions in 10° bins. This prevented event distributions from being confounded by non-uniformities already present in the transform distributions. Resulting circular patterns were visualized through polar histograms (Figure 4E-F).

### Laminar LFP in mice

Z scored physiological fluctuations such as theta frequency were Hilbert-transformed (Figure 5A, left), resulting in curves with one phase value (in degrees) per time sample (at 125 Hz). We then decided to subdivide the infraslow cycles in 45° steps, thus creating eight LFP sub-epochs per cycle. For that, each phase value (one per time sample) was rounded to one of nine phase categories (0°, 45°, 90°, 135°, 180°, 225°, 270°, 315°, 360°), whichever was closest, forming coarse phase curves (Figure 5A, right). Values rounded to 360° were reassigned as 0°, which resulted in the final eight infraslow phase categories (Figure 5B). The boundaries between 45° steps were taken as timestamps to subdivide each LFP channel into eight sub-epochs per infraslow cycle (Figure 5A, right). Sub-epochs shorter in duration than 2.5 s were discarded. All other sub-epochs were analyzed for PSD (built-in *pwelch* function, 3 s Hamming window, 50% overlap) in theta (5-10 Hz, 0.1-Hz bins), gamma (40-100 Hz, log-spaced) and HFO bands (100-160 Hz, log-spaced) or phase-amplitude cross-frequency coupling, CFC (e.g., Brankačk et al., 2012; Scheffer-Teixeira and Tort, 2017), with phase frequencies set to 4-12 Hz (0.25-Hz bins) and amplitude frequencies defined as 20-200 Hz (5-Hz bins).

These methods were used in all analyses of Figure 5C-G. The images in panels C and F of Figure 5 are from averaging across all home-cage sessions from all mice. The curves in panels D, E, G, H of Figure 5 are specific to the physiologically relevant categories 0° and 180° (Figure 5B), with variations (shaded areas) representing the home-cage sessions.

### Laminar firing in mice

#### Spike sorting

Neuronal spikes were extracted from wideband data using Kilosort2 (Stringer et al., 2019), a modification of Kilosort (Pachitariu et al., 2016) that accounts for silicon probe drift, enabling better tracking of single units across long recordings like the ones done in this study. Parameters used were: minimum firing rate per channel = 0.1 Hz, threshold = [10 4], lambda = 10, AUC = 0.9. This was followed by manual curation of putative neuronal units in Phy. Units that dropped out during a recording were excluded from analysis.

#### Spike-phase relationships and firing rates

Spikes were examined for two types of phase relationship: (1) with Z-scored infraslow physiological fluctuations (theta frequency, theta power and EMG from REM and WAKE; Figure 6A-C) and (2) with theta-filtered LFP divided into the coarse infraslow sub-epochs from theta frequency fluctuations, specifically (Figure S5A). In both cases, circular distributions from spike phases were normalized against their underlying transform distributions as done previously (Figure 4E-F). In Figure 6D-E and Figure S5C, phase relationships were represented by scatter plots, not polar histograms, allowing visualization of the multiple neuronal units. Each unit was represented by a data point in the scatter plots, with X axis positions and data point diameters representing mean angles (*circ_mean* function) and mean resultant lengths (MRL; *circ_r* function), respectively (Berens, 2009). Neuronal depths were scattered along the Y axes by assigning units to their anatomically mapped channels (1-64). Channels were slightly jittered by subtracting from each of them a random number from 0–1, allowing better visual assessment of the data. Finally, channels were converted to millimeters given the known electrode spacing along the probe (20 μm). To complement Figure 6D-E, in particular, neurons were classified into putative excitatory (pE) and inhibitory (pI) based on their firing rates and the width of their action potential waveforms (pI: faster firing, narrower waveforms; Stark et al., 2013; Watson et al., 2016) and color-coded accordingly. Then two types of histograms (10° bins) were produced from scatter plot data: (1) the sum of neuronal units across depths followed by probability density function normalization (PDF) and (2) the average MRL across depths.

Spikes were supplementarily examined for their firing rates across coarse infraslow sub-epochs. As depicted in Figure S5B and S5D, we obtained spike counts per infraslow sub-epoch, whose durations were also quantified. Then we sorted both spike counts and sub-epoch durations into the eight infraslow categories previously used in Figures 5-6. The total spike count was divided by the total duration of each category, resulting in firing rates. This was done iteratively across neuronal units, which were binned into channels (1-64) producing firing rate matrices with depths in dimension 1 and the eight infraslow categories in dimension 2. Multiple matrices like these were produced, each one representing a fluctuating physiological signal (theta frequency, theta power and EMG from REM and WAKE). Such matrices were concatenated horizontally, i.e., across physiological signals, normalized from 0–1 per depth, and de-concatenated back into their respective physiological signals, culminating in the heatmaps of Figure S5D. This entire process was done separately for pE and pI units. Finally, we quantified the coarse phase patterns seen in the heatmaps by averaging across depths, and again doing 0–1 normalization across physiological signals, resulting in the phase histograms of Figure S5D.

#### Cell assemblies

Assembly patterns were detected and assembly activity quantified as described in Lopes-dos-Santos et al., 2013. For each recording, episodes of REM or duration-matched episodes of WAKE were concatenated and converted into spike count matrices with 100-ms bins. The spiking activity of each unit was then standardized across time by Z scoring. For each pair of units, (1) the linear correlation was used to populate a correlation matrix for each session and (2) the principal components were extracted. For pairs whose activity is uncorrelated, the variance along each principal component has an upper bound defined by the Marčenko–Pastur distribution (Peyrache et al., 2009). Variances that exceeded this bound were taken to indicate principal components that violated this null hypothesis of independent activity and were therefore statistically significant. This yielded 90 assemblies in REM and 104 assemblies in WAKE across 15 home-cage sessions from 5 mice. The standardized spike matrix was projected onto the subspace defined by the significant principal components to reduce its dimensionality. Then, independent components were extracted via fast ICA (Hyvärinen and Oja, 1997). The independent components attribute large weights to select units, defining that assembly pattern.

Following assembly detection, assembly activity was calculated as the weighted sum in each 100-ms bin of the standardized spike matrix. This temporal resolution allowed us to sub-epoch assembly activity into the eight infraslow categories, as illustrated in Figure 6F. The mean assembly activity in each infraslow category was then plotted to form violin distributions (Figure 6G). Each distribution yielded a mean coactivation strength value (see white circles in Figures 6G and S6). These values were observed to fluctuate across infraslow categories, and we correlated them against sinusoidal fits. Each sinusoidal fit was determined by setting the period equal to that of the infraslow oscillation (nine categories, corresponding to the full 180°–180° cycle), vertical shift equal to the mid-range of the fluctuation, and amplitude equal to half the range. The phase shift was varied from 0 to 2π in 100 increments, and the value that yielded the greatest correlation between the sinusoid and the fluctuation was used.

### Human polysomnography

#### Data collection and sleep staging

All human subjects underwent a Type I, attended, in-lab overnight PSG using Polysmith software (Nihon Kohden, Japan) in an American Academy of Sleep Medicine (AASM)-accredited sleep laboratory at a single site (VAPORHCS). Six scalp electrodes were placed at F3, F4, C3, C4, O1 and O2 per the 10-20 system of EEG placement. Following the conclusion of each recording, an AASM-accredited technician manually scored sleep stages according to AASM-standard clinical criteria (manual version 2.6). Each 30-second epoch of data was scored as one of the five sleep stages: WAKE, REM and NREM stages N1, N2 and N3. Staging of each PSG was additionally validated by a board-certified sleep physician.

#### Post-curation PSG analysis

EEG channels were upsampled from their original digitization rate (200 Hz) to the raw LFP rate of the mouse portion of this study (1250 Hz) using linear interpolation. This made the PSG data compatible with the same sleep scoring spectrogram methods mentioned earlier (1-s bins, 1–40-Hz linearly spaced frequencies). The exception is that we did not perform the sleep scoring on the raw data. Instead, we adapted the existing 30-s-binned states assigned by the PSG technician to the 1-s binning of our spectrograms, and then manually adjusted REM epochs to include 5–10-min margins for later refinement at 125 Hz, similar to what we did in mice (compare Figures 1D and 7B).

WAKE epochs from the data curation and sleep staging methods explained above required further elimination of periods contaminated by movement artifacts. In addition, total WAKE time was lower than REM time in all human hypnograms we analyzed, as opposed to our mouse hypnograms, whose recordings were more abundant in total WAKE time. Thus, in humans we could not perform the duration-matched REM-WAKE epoch pairing approach we did in mice. However, this was not detrimental to our usage of human WAKE as control in infraslow periodicity analysis (see below) because all human WAKE and REM epochs that we approved for final analysis were longer than 5 min (i.e., longer than in mice; Figure S1D). NREM epochs were also obtained per human subject for supplementary REM vs. NREM comparisons (Figures S7 and S8). See Table S2 for number and duration of epochs from the entire human dataset.

From the epochs described above, we analyzed the incidence of rapid movements vs. periods of motoric quiescence, as well as variations in respiration and heart rates and band power curves. Methods were similar to those used in mice (compare Figure 7B with Figures 2B and 3C-E). Specifics of human data are explained as follows. EOG was processed by calculating the absolute difference between left and right EOG signals, generating zero-based curves containing spike-like events. Alternating periods with low and high incidence of such events were captured using peak envelopes (10-s window).

The two leg signals, left and right, were averaged to produce one combined signal. We then analyzed the incidence of leg movements using the same peak envelopes as above. Respiration and heart rates were resampled to 125 Hz and analyzed using wavelet spectrograms with linearly spaced frequencies. The respiration rate band was the same across subjects: 0.1–0.5 Hz (0.002-Hz bins). Heart rates, on the other hand, were more prone to individual variability, and therefore their bands were adjusted per human subject. The slowest and fastest heart rate bands we used were, respectively, 0.6–1.4 Hz and 1.3– 2.0 Hz (0.005-Hz bins regardless of the band). All respiration and heart rate spectrograms showed robust frequency-varying trails irrespective of brain state (Figures 7B and S7). Hence, each spectrogram was analyzed using a single time-frequency ridge, without necessity of multiple ridges as we did in mice (Figure 3D). Finally, band power curves from four bands (human theta: 4-8 Hz, alpha: 8-12 Hz, beta 1: 12-20 Hz, beta 2: 20–30 Hz) were obtained by averaging across the frequency bins of EEG spectrograms, also at 125-Hz sampling.

We then proceeded to three types of quantitative analyses: distribution bimodality vs. unimodality (histograms: 100 bins; Figure 7C-D), infraslow periodicity (built-in *pwelch* function, 5-min Hamming window, 50% overlap, 0–0.1-Hz band, 0.001-Hz bin size) in REM vs. WAKE (Figure 7E) and phase relationship between movement events and infraslow physiological fluctuations (polar histograms: 10° bins; Figure 7F). All of these analyses were considered optimal in dissecting REM microstructure based on what we learned from the mouse portion of this study, and thus we decided to use them for the human data. Key differences between human and mouse analyses were adaptations to two parameters: the envelope window to examine movement bimodality/unimodality (10 s in humans, 2 s in mice) and the infraslow periodicity window (5 min in humans, 3 min in mice). These parameters were determined heuristically and reflect the different features of intra-REM fluctuations in the two species, with both movements and infraslow fluctuations being slower in humans than mice. Finally, EEG montages available in our PSG dataset were averaged into frontal (from F4A1 and F3A2), central (from C4A1 and C3A2) and occipital (from O2A1 and O1A2) signals and had their distributions, infraslow periodicity and phase relationships with movements quantified as above.

### Statistical analysis

In both mice and humans, the distributions of enveloped eye and facial movements (Figures 2C and 7C) were examined for bimodality vs. unimodality using Hartigan’s dip test. The same was done for distributions from physiological fluctuations, which were additionally examined for REM vs. WAKE differences in variance using the two-sampled F-test for equal variances (*vartest2* function; Figures 3G-H, 7D and S3D-E). To compare infraslow PSD curves from REM and WAKE, we used two-way ANOVA (with frequency bins as repeated measures) followed by Tukey’s post-hoc tests at individual frequency bins (built-in *fitrm, ranova* and *multcompare* functions; Figures 3G-H, 7E and S3F-G). Finally, circular statistics consisted of Rayleigh tests (*circ_rtest* function, Berens, 2009) with Bonferroni correction of the alpha value, i.e., 0.05 divided by the number of tests shown in Figures 4E-F, 7F and S8A.

Regarding mouse-specific analyses, the alpha value for the correlation matrices in Figure 4A was also Bonferroni-corrected (number of correlation coefficients from both REM and WAKE: 30). Significant P values were then divided into quartiles and indicated by asterisks on top of the correlation matrices for visual appreciation of significance levels (Figure 4A). The facial quiescence and facial movements clusters in Figure 4B were compared in Figure 4C using two-way ANOVA (facial movement cluster vs. electrophysiological measure; *fitrm, ranova* and *multcompare* functions). Shaded curve graphs across Figure 5 represent several comparisons between infraslow categories 0° and 180°, all of which were made along the depth dimension. These were all quantified using two-way ANOVA (with depths as repeated measures) followed by Tukey’s post-hoc tests at individual depths (built-in *fitrm, ranova* and *multcompare* functions). Rayleigh tests (*circ_rtest* function, Berens, 2009) were performed on all histograms of Figures 6D-E and S5, also with Bonferroni correction of the significance level. Finally, correlation coefficients were obtained between mean assembly activity in each infraslow category and the sinusoidal fits in Figures 6G and S6. Bonferroni-corrected P values are represented by asterisks or groups of asterisks in such figures.

## Supplemental videos

**Video S1. Example head-fixed REM episode with facial movements and brain electrophysiology shown in 1x time speed (related to Figure 2)**.

**Video S2. Laminar LFP patterns along the infraslow cycle of REM theta frequency fluctuations (related to Figure 5)**.

---

**Figure S1.**
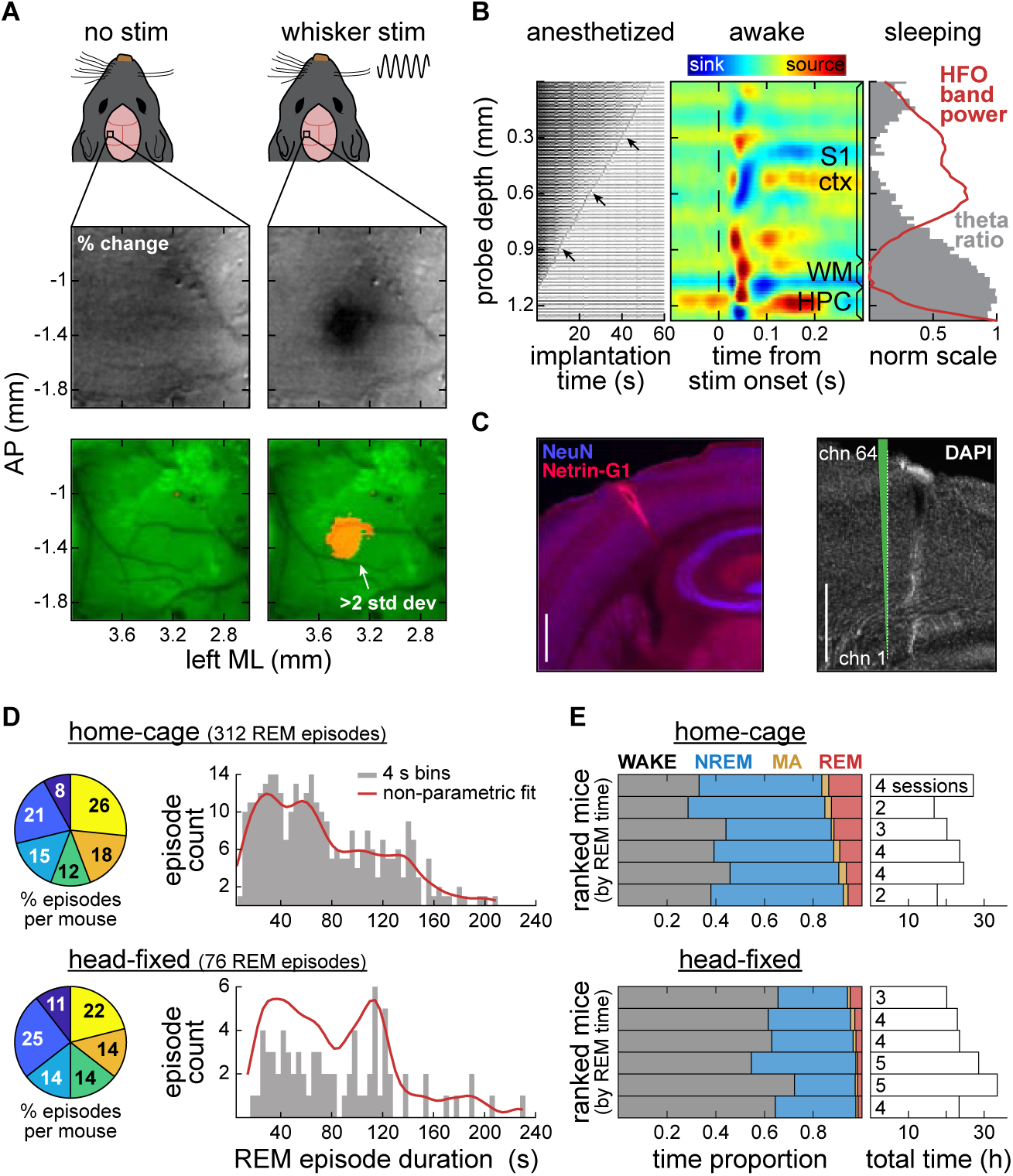
Targeted silicon probe implantation and incidence of REM episodes across mice (related to Figure 1). (A) Intrinsic optical imaging of primary somatosensory cortex (S1) under anesthesia. Top: vibrotactile stimulation (stim) and skull thinning above the D1-2 barrel vicinity. Center: average percent change images from stim and no-stim trials revealing barrels via blood oxygen signals (Petersen, 2007). Barrel response is seen here as the darkened area in the center of stim image. Anteroposterior (AP) and mediolateral (ML) coordinates are based on a canonical map of the barrel field (Knutsen et al., 2016). Bottom: statistically significant change during stim versus no-stim control (orange). By superimposing the barrel signal on a blood vessel image (green), we produced mouse-specific implantation targets. (B) Electrophysiology-guided silicon probe lowering under anesthesia using a micromanipulator. Left: representative raw recordings from the 64 channels during probe lowering into S1. Distance between electrode sites: 20 μm. Voltage transitions from noise to brain signal occurred as electrode sites first entered the brain through the pierced dura mater (see arrows). We ceased the dorsal-ventral trajectory once the most superficial channel completed the transition from noise to signal. Center: post-surgery somatotopic validation under awake head fixation (separate from the main sleep recordings of this study). Current source density (CSD) around whisker deflection stimuli (0.2-s duration) suggested a depth profile reaching beyond S1 cortex (S1 ctx) into the underlying white matter (WM) and hippocampus (HPC). The whisker pair yielding maximal CSD responses was kept untrimmed for the remainder of the study. This was intentionally redundant with the procedure in (A) to provide extra targeting. Right: additional laminar profiling using per-channel theta ratio and band power of high-frequency oscillations (HFO). (C) Histology confirming the depth profile determined physiologically. Left: merged NeuN and Netrin-G1 immunofluorescence highlighting the neocortical laminar structure and the hippocampal pyramidal layer, both of which were sampled by the probe. Implantation angle: 30°. Right: rotated DAPI immunofluorescence showing the probe tract in parallel to a depiction of the probe (chn: channel). Both scale bars: 0.5 mm. (D) Left: distribution of home-cage and head-fixed REM episodes across mice (n = 6). One color per mouse in each pie chart. All mice contributed in a balanced manner to both REM episode counts even though the incidence of REM was lower under head fixation as expected. Right: distribution of REM episode durations (4-s bins). The home-cage dataset formed a multimodal distribution of REM episode durations, reinforced by the multiple peaks of a non-parametric fit (kernel smoothing window: 8 s). We did not explore this finding further in this study. A similar distribution could not be replicated in the head-fixed dataset, possibly due to its lower number of REM episodes. (E) Proportional time spent in each brain state, including microarousals (MA). Left: each row represents a mouse. The higher the row position, the higher the proportional time spent in REM for that mouse. Sleep states were generally more difficult to reach under head fixation, consistent with the lower number of head-fixed REM episodes. Right: amount of time (bars) and sessions (numbers inside bars) contributed by each mouse (same ranking as the left plots). Comparable with Tsunematsu et al. (2020).

**Figure S2.**
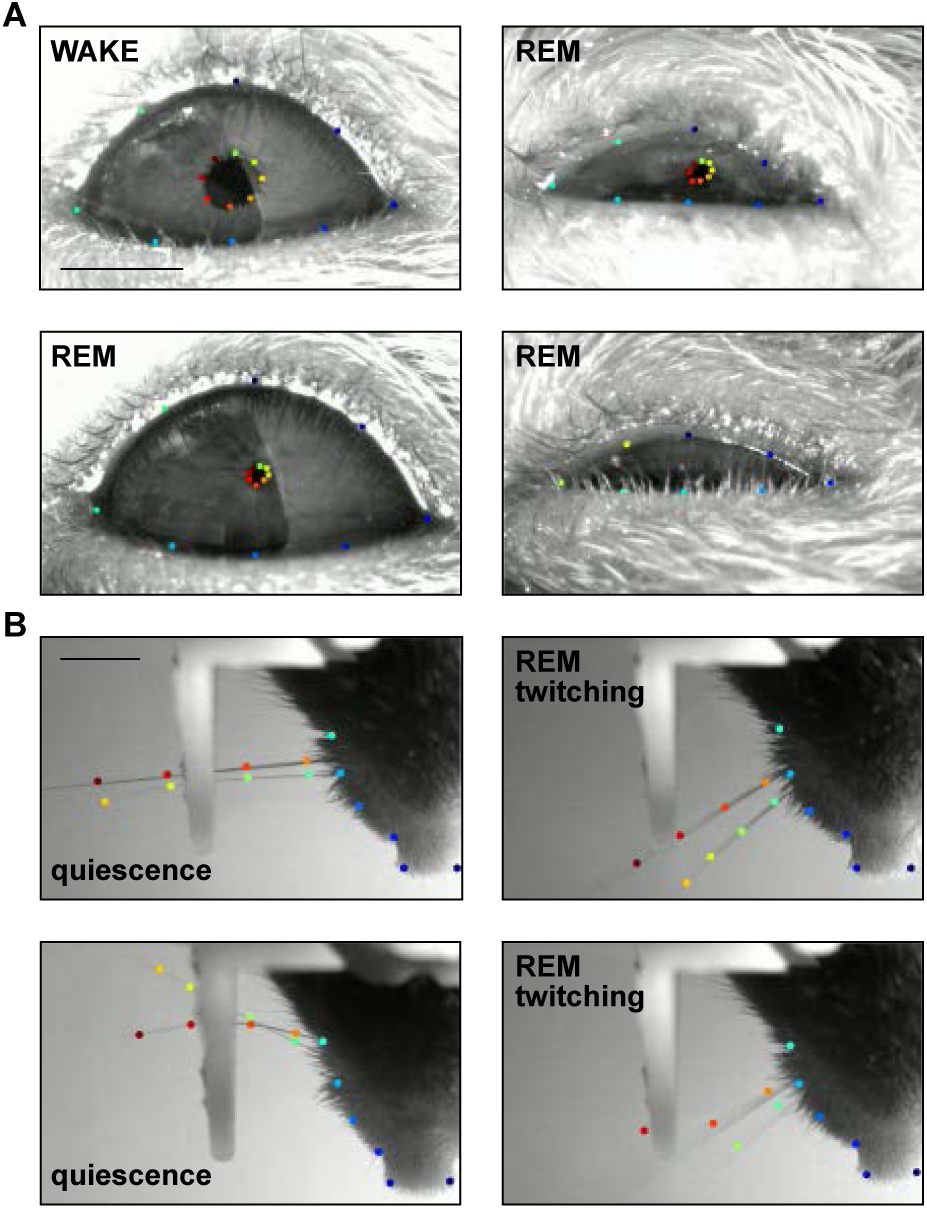
Representative DeepLabCut-labeled frames from mouse REM and WAKE (related to Figure 2). (A) Typical eye video scenarios and corresponding octagonal labeling of the eyelids and pupil. The top-left frame shows the eyelid separation and pupil size normally seen during quiet WAKE. All other frames were collected during REM, characterized by constricted pupil and varying eyelid state, from wide open (bottom-left frame) to almost closed (right frames). The bottom-right frame exemplifies a hidden pupil situation, from which we could still record the myoclonic eye blinks that generally accompanied rapid eye movements (see also Table S1). Incomplete eyelid closure during sleep in mice was also reported by Yüzgeç et al. (2018) and Sattler and Wehr (2021). Scale bar: 1 mm. Recording rate: 20 frames/s. (B) Typical whisker frames and corresponding labeling on whiskers, mystacial pad and nose. The left frames show pairs of untrimmed whiskers in rest position, observable either in the quiescent periods of REM and WAKE or across NREM. The right frames represent the apex of REM twitches. In a minority of frames with motion blur and erratic labeling (bottom right), whisking could still be measured with little impact since whisker speed was obtained by averaging across whisker labels for redundancy. This approach, in addition to our recording rate (100 frames/s, lower than whisker-focused studies) was sufficient to capture bursts of REM twitches. Scale bar: 3 mm.

**Figure S3.**
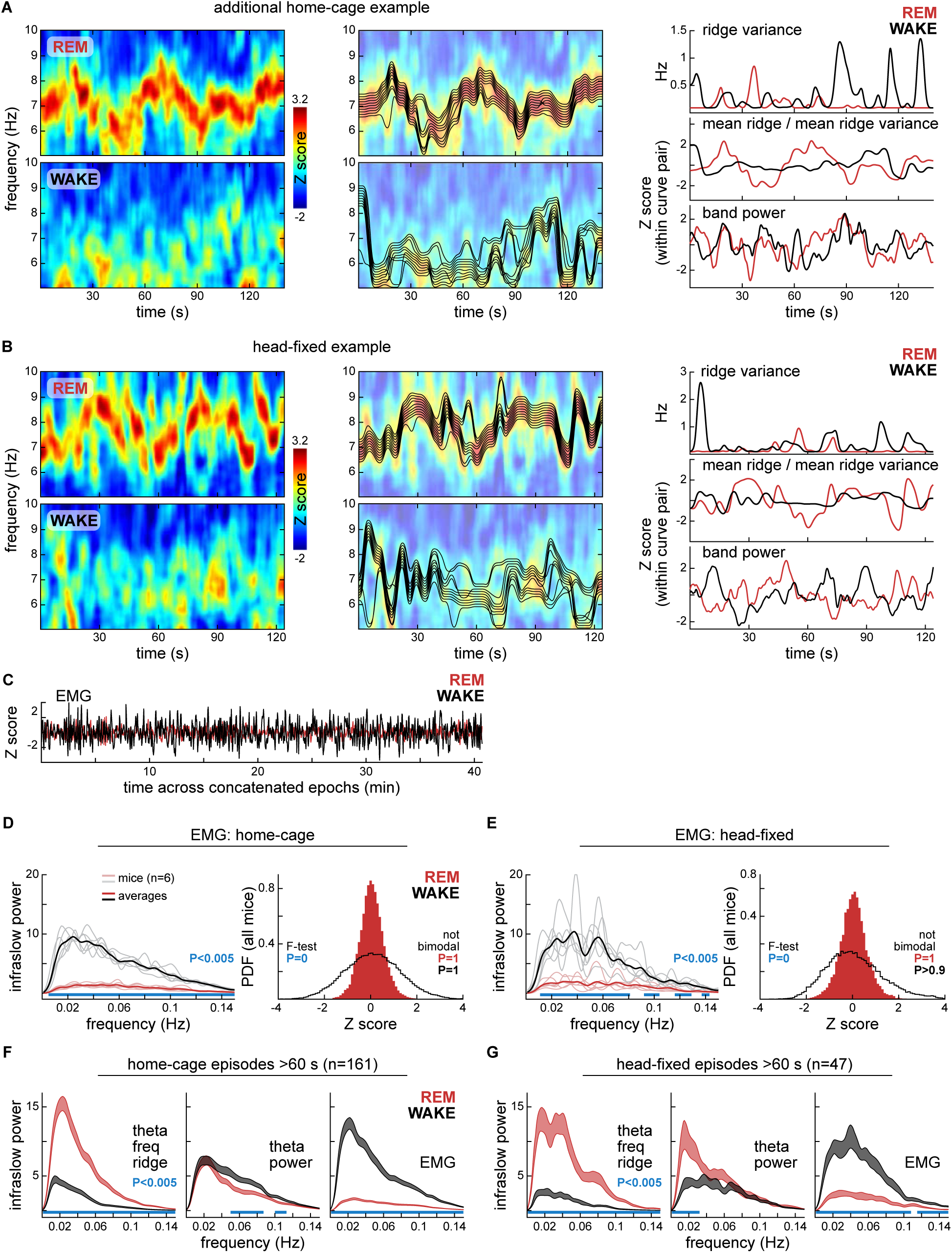
Additional spectrogram examples and additional controls for infraslow PSD analysis in mice (related to Figure 3). (A) Additional example from the home-cage dataset. See full explanation in Figure 3. (B) Example from the head-fixed dataset. (C) EMG epochs from REM or WAKE were concatenated as in Figure 3F. We again observed minute-scale fluctuations, but with higher amplitude in WAKE. (D) Spectral content and distribution of concatenated EMG curves. Data from all home-cage sessions are shown (312 REM-WAKE epoch pairs, 6 mice), as in Figure 3G. Left: infraslow power spectral density (PSD) per mouse (faded lines) and averaged across mice (bold lines). Blue bars at bottom indicate post-hoc differences at individual frequency bins (bin size: 0.001 Hz) after two-way ANOVA (state vs. frequency). Infraslow rhythms were stronger in WAKE EMG (interaction: F_1,10_ = 650.09, P = 1.98*10^−10^). Right: sample count histograms from all sessions. We again found no evidence of bimodality (Hartigan’s dip test). Variance was greater in WAKE (F-test of equality of variances; F = 0.18, P = 0). (E) As in (D), but from head-fixed data (76 REM-WAKE epoch pairs, 6 mice; compare with Figure 3H). ANOVA interaction: F_1,10_ = 71.95, P = 7.03*10^−6^. F-test of equality of variances; F = 0.24, P = 0 (F) Infraslow spectra from REM and WAKE home-cage epochs longer than 60 s (161 of 312 epoch pairs). Epochs were examined for infraslow PSD without concatenation as an additional control for Figure 3G. Shaded curves delimit standard errors from the 161 epochs. All spectra confirmed those obtained with the concatenation of theta signals (Figure 3G) and EMG (panel D of this figure). Blue bars at bottom indicate post-hoc differences at individual frequency bins (bin size: 0.001 Hz) after two-way ANOVA (state vs. frequency). Theta frequency effect of interaction: F_1,10_ = 159.31, P = 6.46*10^−30^. (G) As in (F), but from long-duration head-fixed epochs (47 of 76 epoch pairs). Compare with Figure 3H and panel E of this figure. Theta frequency effect of interaction: F_1,10_ = 142.21, P = 2.27*10^−20^.

**Figure S4.**
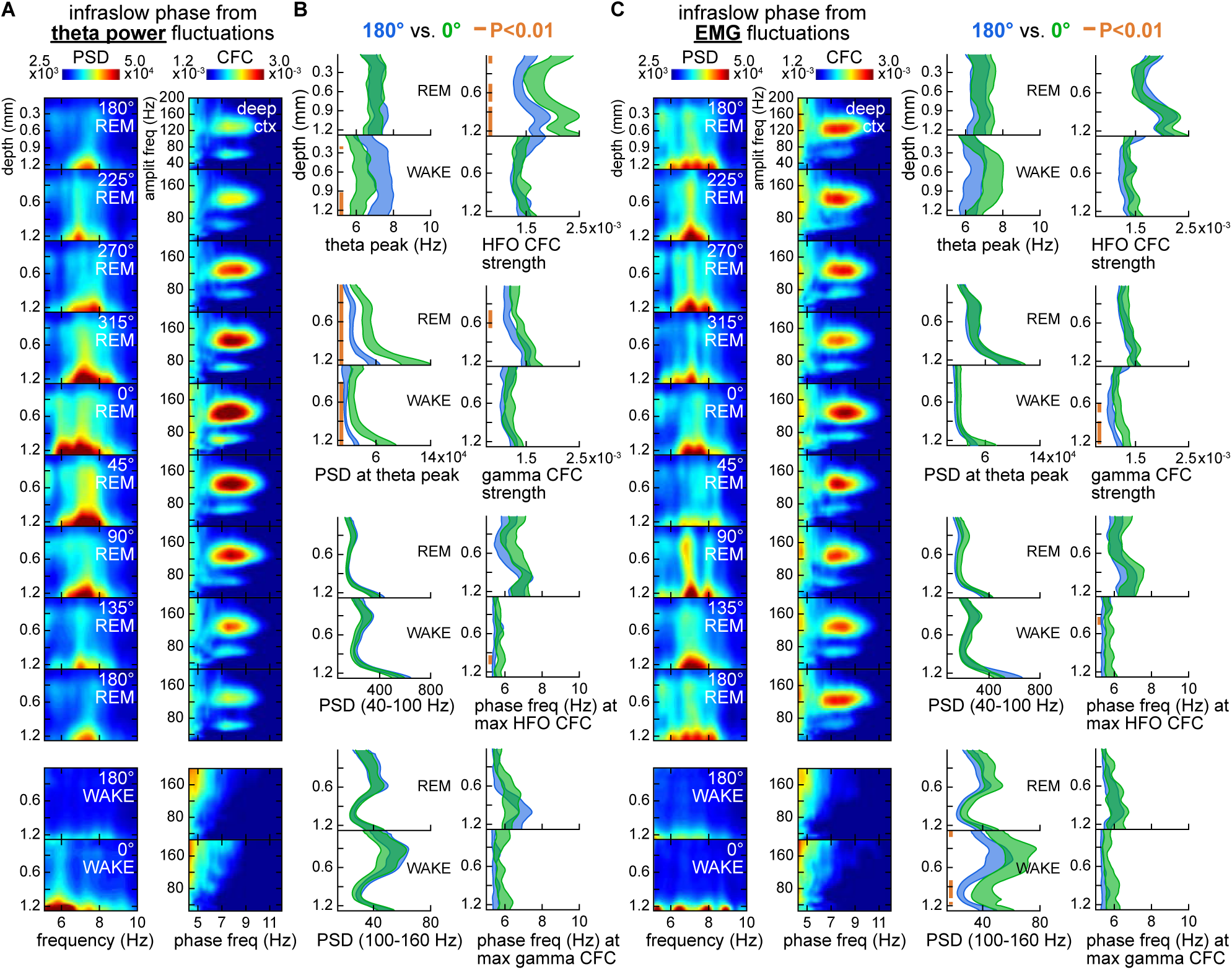
Laminar LFP by the infraslow phase of theta power and EMG in mice: additional controls for infraslow sub-epoching (related to Figure 5). (A) Laminar LFP by the coarse infraslow categories from theta power fluctuations (see method in Figure 5A-B). Left column of subplots: power spectral density curves (PSD, color axis) along theta frequencies (5–10 Hz, X axes) were stacked across channels (Y axes) then averaged across mice (home-cage sessions only), constructing one laminar profile per infraslow phase (vertical sequence of subplots), similar to Figure 5C. PSD was stronger closer to the 0° infraslow category as expected. In this analysis, 0° is not necessarily coincident with the “phasic” REM pole explained in Figure 5B since according to Figure 4 the infraslow fluctuations from theta power were not optimally related to REM facial movements. In fact, in this panel the blobs are more dispersed along the theta frequency dimension (X axes), reinforcing that theta power and theta frequency do not necessarily fluctuate together within REM (compare with the more organized blobs of Figure 5C). Right column of subplots: cross-frequency coupling (CFC) between theta (phase frequencies: 4–12 Hz) and faster oscillations (amplitude frequencies: 20–200 Hz) per infraslow category from theta power, similar to Figure 5E. For simplicity, only one representative depth is shown: deep S1 cortex. CFC was stronger closer to the 0° infraslow phase, especially in the high-frequency oscillation blob (HFO, 100– 160 Hz). However, unlike Figure 5E, blobs did not shift horizontally along the theta phase frequency dimension (X axes). Thus, HFO CFC strength may depend on both theta power and theta frequency during REM. WAKE again showed much weaker CFC patterns. (B) Left column of subplots: 0° vs. 180° comparisons within REM or WAKE laminar PSD, corresponding to the left heatmaps of (A). Similar layout and quantification as Figure 5D–E. PSD at theta peak was expectedly stronger in the 0° infraslow category in both REM and WAKE. In WAKE specifically, we also observed that the 0° infraslow category was associated with lower theta peak frequency, especially at deeper channels (see first pair of graphs). We did not explore this finding further. Right column of subplots: 0° vs. 180° comparisons within REM and WAKE laminar CFC. Graphs correspond to the right heatmaps of (A), but from the full laminar profile. Similar layout and quantification as Figure 5G-H. The most evident effect was that the 0° infraslow category of REM showed stronger HFO CFC at most depths, suggesting, along with Figure 5G-H, that laminar HFO CFC varies with the infraslow fluctuation of both theta power and theta frequency in REM. (C) As in (A-B), but with sub-epoching based on the infraslow fluctuation from EMG. Blobs in laminar PSD images are even less organized (see heatmaps on the leftmost column), reinforcing that theta frequency is optimal in determining REM microstructure in mice (Figure 5). HFO CFC did not vary with the EMG infraslow category, both in terms of CFC strength and the phase frequency dimension (see second column of heatmaps and right column of graphs). Laminar HFO power was, however, stronger in WAKE (bottom graph of third column). We also did not explore this observation further.

**Figure S5.**
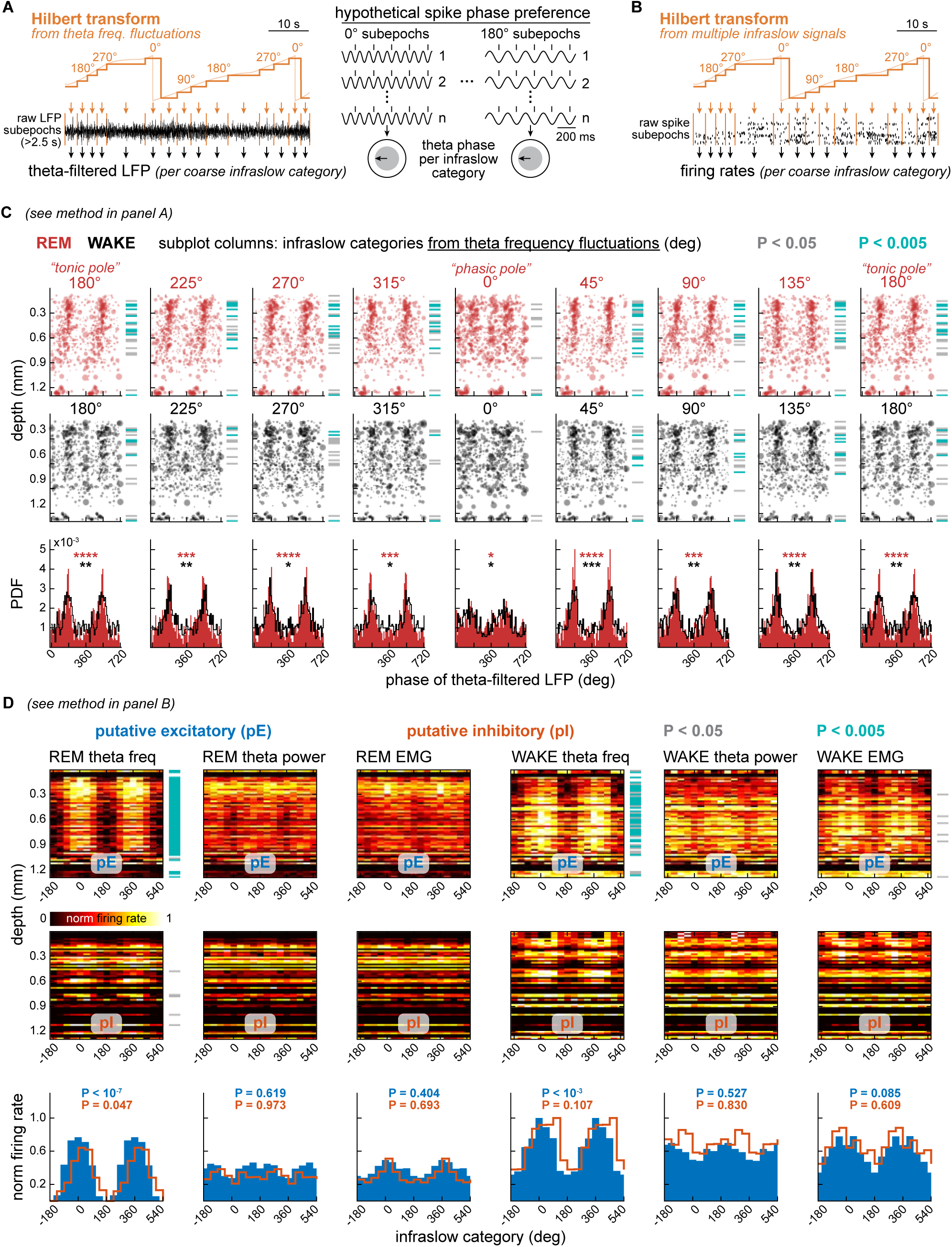
Laminar spiking along the infraslow microstructure of mouse REM: additional evidence that firing rates vary along a non-binary continuum between slow and fast theta activity (related to Figure 6). (A) Left: the same method as in Figure 5A was used to classify sub-epochs of theta-filtered LFP into coarse infraslow categories, specifically from theta frequency fluctuations. Right: each neuronal unit had its phase relationship with filtered theta cumulatively measured across the sub-epochs of each infraslow category. This resulted in theta-phase angles (arrows inside gray circles) and mean resultant lengths (MRL, diameter of gray circles) per category. Circles like these were parsed into anatomical depths, theta phases and infraslow categories, as shown in the scatter plots of (C). (B) Another method to analyze laminar firing across infraslow cycles. Spikes were counted per coarse infraslow category. Spike counts were then divided by the total time of each infraslow category, yielding firing rates. Such firing rates are shown along anatomical depths in the heatmaps of (D). (C) Data from all home-cage sessions from all mice. Scatter plots: infraslow categories are laid out as subplot columns, each column containing REM and WAKE scatter plots, as well as depth-mapped color-coded P values from phase occupancy (Rayleigh test), similar to Figure 6D-E. Histograms: theta phase occupancy across depths (10° bins, probability density function normalization, or PDF), corresponding to the scatter plots. Both scatter plots and histograms show that phase relationships with filtered theta were robust across infraslow categories, except 0°, whose circular patterns were less organized. This could be due to whisking temporarily disrupting the phase relationship between S1 spiking and ongoing theta activity, since according to Figure 4E-F, whisking was more likely around the 0° poles of theta frequency fluctuations in both REM and WAKE. We addressed this potential confound in (D). (D) Data from all home-cage sessions from all mice. Heatmaps: REM and WAKE firing rates per anatomical depth (Y axes) and coarse infraslow category (45° bins, X axes) from all infraslow fluctuations (subplot columns). Additional distinction is made between putative excitatory and inhibitory neurons. Histograms: firing rate across depths per coarse infraslow category (45° bins, normalized on a 0–1 scale), corresponding to the heatmaps. REM theta frequency showed the most significant circular patterns, consistent with Figure 6D-E. Although strong, these circular patterns were not dominated by the 0° category. This, along with panel (C), suggests that ongoing theta activity — not whisking — is the main driver of the results in Figure 6D-E, reinforcing the continuous nature of intra-REM fluctuations other than facial motor activity.

**Figure S6.**
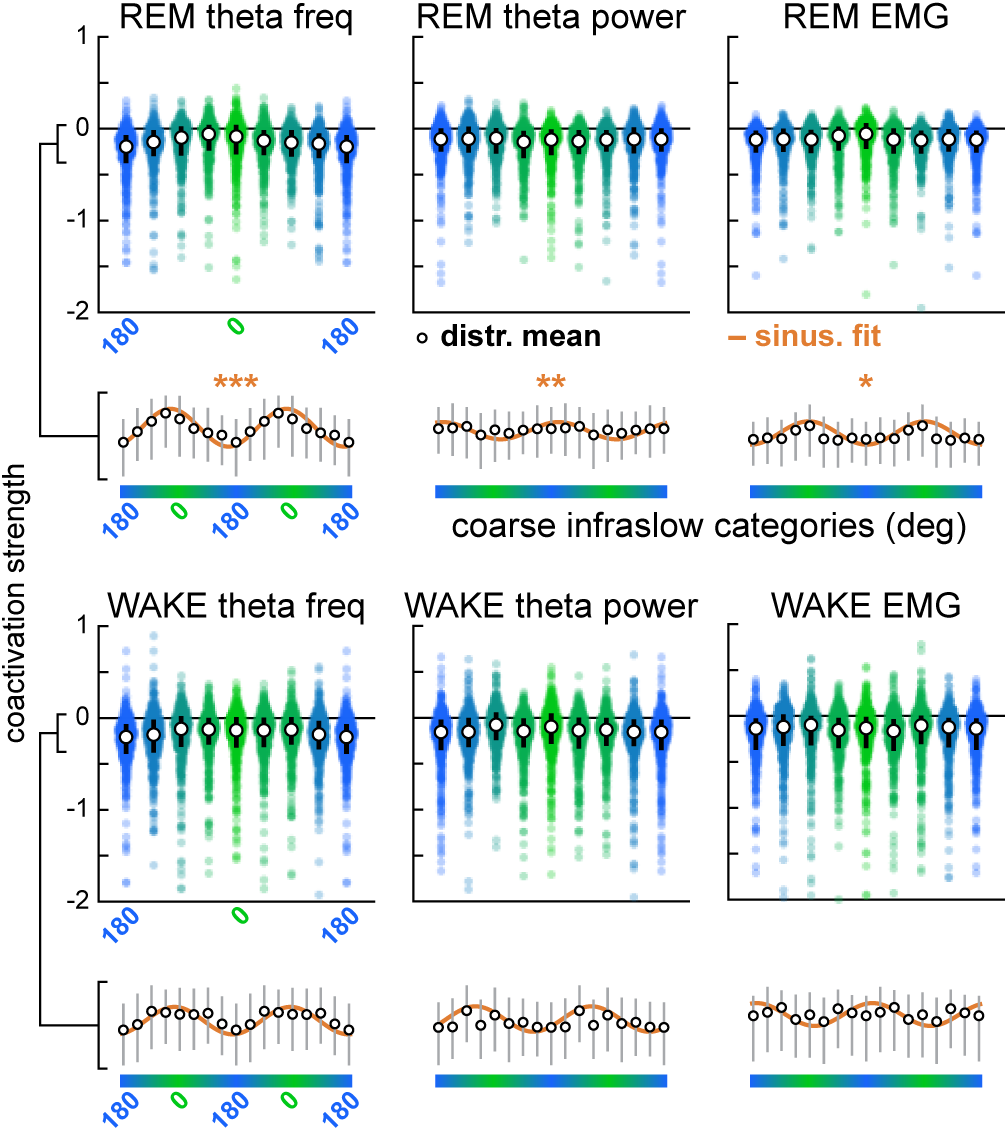
Alternative assembly analysis method corroborates REM theta frequency as an important driver of spiking coordination (related to Figure 6). Similar to Figure 6G. However, for this figure, Hebbian-inspired ICA-based cell assemblies (Lopes-dos-Santos et al., 2013) were detected within each infraslow category rather than within all of REM or WAKE. This yielded a corresponding increase in the number of assemblies detected per session, roughly eightfold. So that coactivation strengths could be compared across assemblies detected in different categories, strength values in each category were normalized as a log fold change relative to the value in the category in which that assembly was detected. Results are essentially the same as in Figure 6G: the fluctuation in coactivation strength across infraslow categories was most sinusoidal relative to the oscillation of REM theta frequency, as quantified by Pearson’s correlations between sinusoidal fits and mean coactivation strengths (*P < 10^−2^, **P < 10^−3^, ***P < 10^−6^).

**Figure S7.**
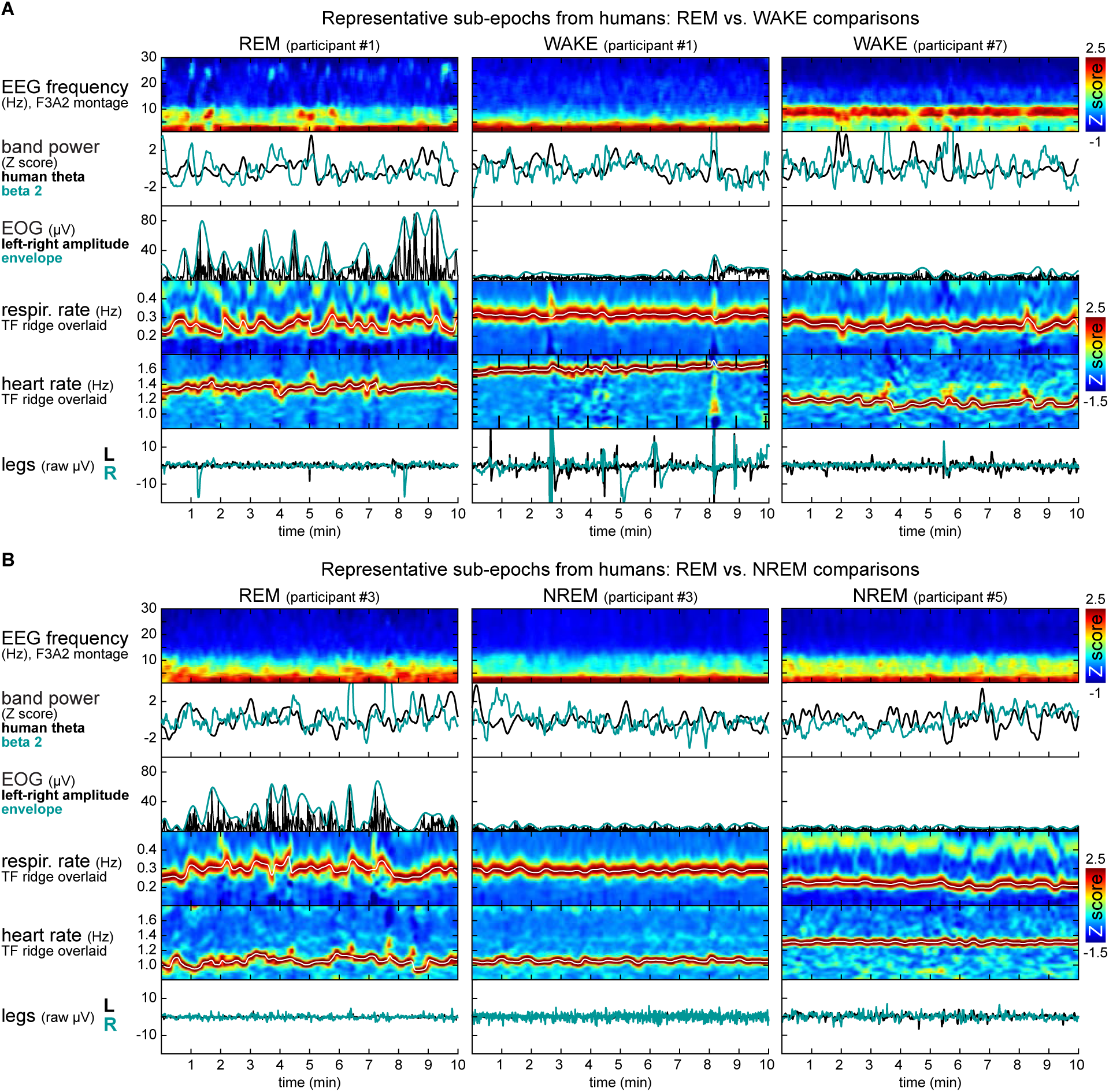
Additional representative sub-epochs from human REM: qualitative comparisons with WAKE and NREM (related to Figure 7). (A) REM and WAKE physiological measures are qualitatively compared within the same participant (left and center). An extra WAKE sub-epoch from a different participant is shown on the right, illustrating variations in WAKE. In these WAKE examples, we show higher incidence of leg movements in participant #1 and higher alpha power (8-12 Hz) in participant #7, possibly reflecting different levels of relaxation and/or different signal qualities. Despite variations like these, which we did not investigate further, WAKE was consistently more stable than REM in both EOG activity and respiration rate, which according to Figure 7 are optimal markers of intra-REM rhythms in humans. All representative sub-epochs were extracted from larger epochs and trimmed to a common duration (10 min) for visual appreciation of minute-scale fluctuations, unlike Figure 7B. (B) As in (A) but comparing REM and NREM. NREM was much more stable than REM in respiration and heart rates despite variations in their baseline levels across individuals. This reinforces the uniqueness of REM in terms of minute-scale physiological rhythms. See Figure S8B for quantitative REM-NREM comparisons.

**Figure S8.**
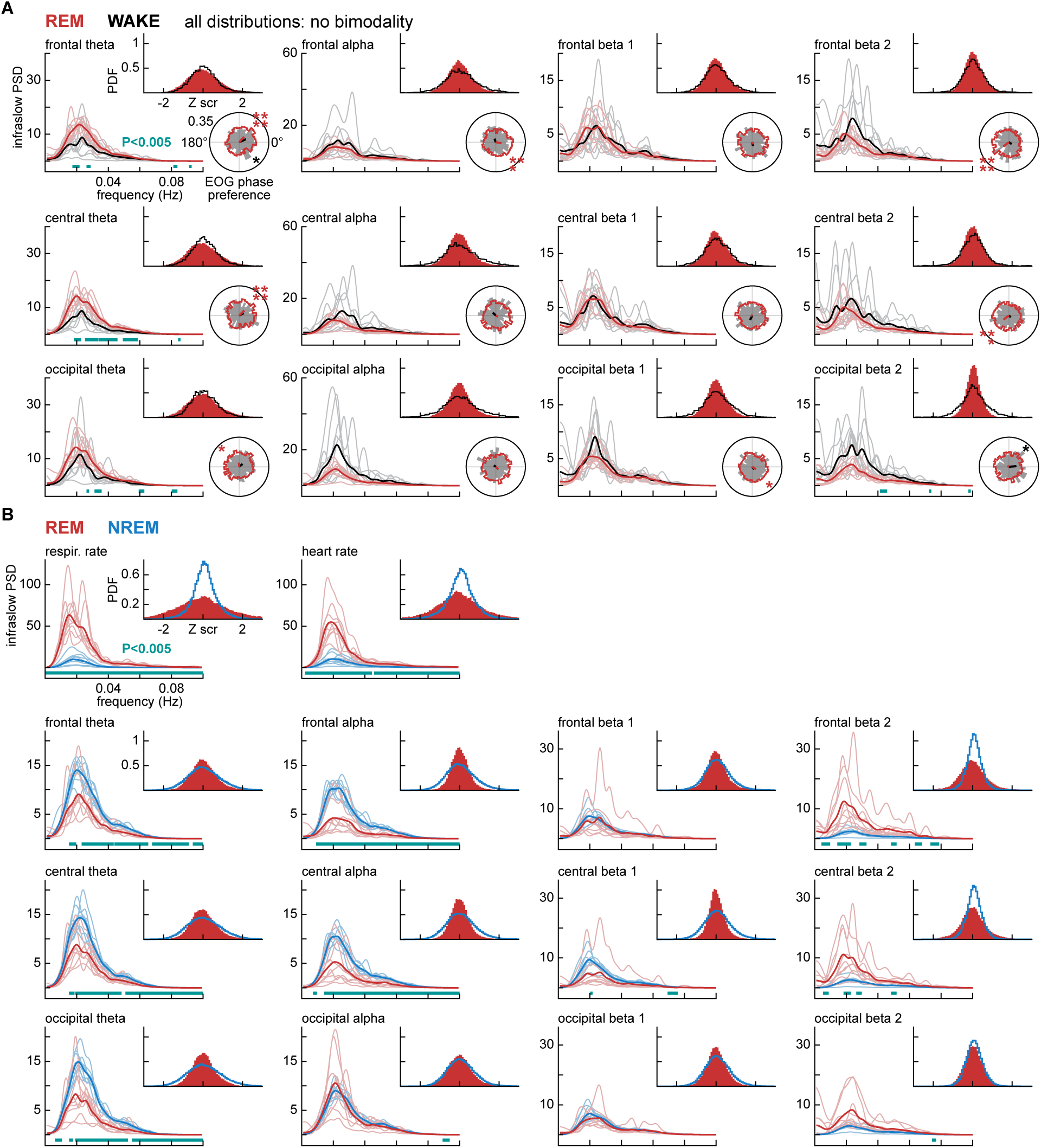
Additional evidence of continuous infraslow fluctuations in human REM: quantitative comparisons with WAKE and NREM (related to Figure 7). (A) Each subplot shows probability density function (PDF) distributions and infraslow PSD from different band power curves (subplot columns) at broad neuroanatomical areas (subplot rows), as well as phase relationships between these band power curves and EOG events (polar histograms). No bimodality was found across distributions, and infraslow spectra did not show evident variation across neuroanatomical areas. This suggests that the nature of band power fluctuations in REM - i.e., continuous and infraslow - is neuroanatomically nonspecific, at least in terms of the EEG montages we analyzed here (mastoid-referenced frontal, central and occipital). EOG phase relationships, on the other hand, did vary across montages, suggesting that band power fluctuations across neuroanatomical areas are not jointly aligned with eye movement density. Data from frontal oscillations are the same as Figure 7. (B) Similar to (A) but comparing REM and NREM. Infraslow fluctuations in respiration and heart rate are much stronger in REM than NREM, reflecting the stability of these physiological measures in NREM (see also Figure S7B). In contrast, REM and NREM showed varying relative strengths of infraslow PSD across the analyzed bands, with NREM being generally stronger in theta and alpha fluctuations (comparable with: Lecci et al., 2017; Osorio-Forero et al., 2021) and REM being stronger in beta 2 fluctuations (consistent with beta 2 variation in human REM: Simor et al., 2020). EOG phase relationships were not analyzed due to the lack of eye movements in NREM.

**Table S1.**
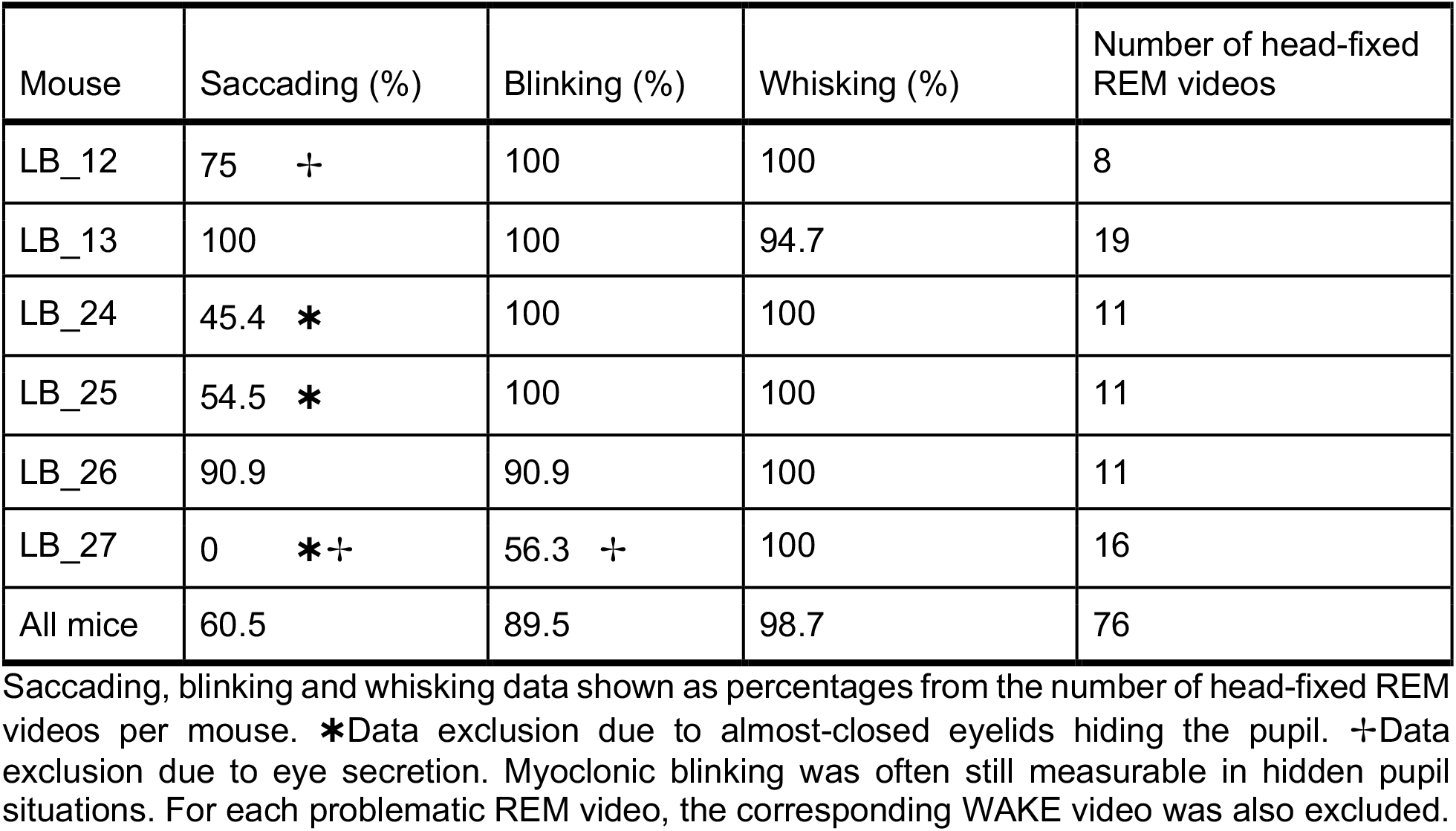
Yield of video data and main reasons for video exclusion (related to Figure 2).

| Mouse | Saccading (%) | Blinking (%) | Whisking (%) | Number of head-fixed REM videos |
| --- | --- | --- | --- | --- |
| LB_12 | 75 † | 100 | 100 | 8 |
| LB_13 | 100 | 100 | 94.7 | 19 |
| LB_24 | 45.4 * | 100 | 100 | 11 |
| LB_25 | 54.5 * | 100 | 100 | 11 |
| LB_26 | 90.9 | 90.9 | 100 | 11 |
| LB_27 | 0 *† | 56.3 † | 100 | 16 |
| All mice | 60.5 | 89.5 | 98.7 | 76 |

Saccading, blinking and whisking data shown as percentages from the number of head-fixed REM videos per mouse. \*Data exclusion due to almost-closed eyelids hiding the pupil. †Data exclusion due to eye secretion. Myoclonic blinking was often still measurable in hidden pupil situations. For each problematic REM video, the corresponding WAKE video was also excluded.

**Table S2.**
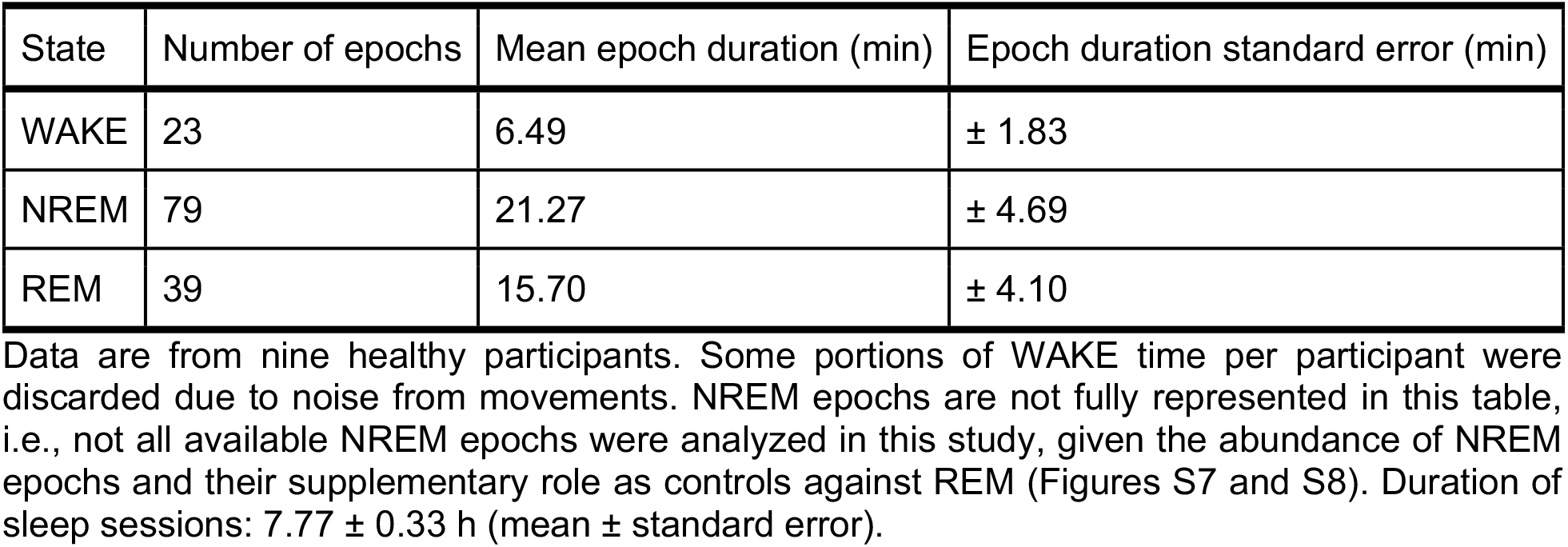
Human polysomnography: number of epochs per state and their duration (related to Figure 7).

| State | Number of epochs | Mean epoch duration (min) | Epoch duration standard error (min) |
| --- | --- | --- | --- |
| WAKE | 23 | 6.49 | $\pm 1.83$ |
| NREM | 79 | 21.27 | $\pm 4.69$ |
| REM | 39 | 15.70 | $\pm 4.10$ |

